# Vocal learning-associated convergent evolution in mammalian proteins and regulatory elements

**DOI:** 10.1101/2022.12.17.520895

**Authors:** Morgan E. Wirthlin, Tobias A. Schmid, Julie E. Elie, Xiaomeng Zhang, Varvara A. Shvareva, Ashley Rakuljic, Maria B. Ji, Ninad S. Bhat, Irene M. Kaplow, Daniel E. Schäffer, Alyssa J. Lawler, Siddharth Annaldasula, Byungkook Lim, Eiman Azim, Zoonomia Consortium, Wynn K. Meyer, Michael M. Yartsev, Andreas R. Pfenning

## Abstract

Vocal learning, the ability to modify vocal behavior based on experience, is a convergently evolved trait in birds and mammals. To identify genomic elements associated with vocal learning, we integrated new experiments conducted in the brain of the Egyptian fruit bat with analyses of the genomes of 222 placental mammals. We first identified an anatomically specialized region of the bat motor cortex containing direct monosynaptic projections to laryngeal motoneurons. Using wireless neural recordings of this brain region in freely vocalizing bats, we verified that single neuron activity in this region relates to vocal production. We profiled the open chromatin of this vocal-motor region, which we used to train machine learning models to identify enhancers associated with vocal learning across mammals. We found 201 proteins and 45 candidate enhancers that display convergent evolution associated with vocal learning, many of which overlapped loci associated with human speech disability. One such locus contains the neurodevelopmental transcription factors *TSHZ3* and *ZNF536* and multiple candidate vocal learning-associated enhancers, suggesting the co-evolution of protein and regulatory sequences underlying vocal learning.

**One-Sentence Summary:** Analyses of bat neural activity and epigenomic data in a brain region involved in vocal behavior were used to identify proteins and regulatory elements associated with vocal learning in mammals.

## Main Text

Vocal learning—the ability of an organism to modify its vocal output as a result of social and acoustic experience—is an example of convergent evolution, having evolved independently within multiple lineages of birds and mammals, including humans, where it manifests as speech (Fig. 1A) (*1, 2*). Vocal learning has been extensively studied in songbirds, highlighting numerous shared behavioral features of birdsong and speech learning, including a dependence on auditory input during a critical developmental period and a juvenile babbling phase of sensorimotor exploration prior to the maturation of the adult repertoire among other features (*1*). Intriguingly, convergence between song-learning birds and humans extends to brain anatomical specializations as well, including direct corticospinal projections from the vocal motor cortex analog to the hindbrain region controlling the vocal apparatus (*3*) and shared transcriptional specializations in analogous speech- and song-specialized brain regions (*4*). Songbirds have thus become a premier model for exploring the fundamental brain anatomical, molecular, and genomic features associated with vocal learning (*1, 3*). An expanding literature on vocal learning behavior across mammals suggests an underappreciated diversity in the phenotypic expression of vocal learning across the taxa traditionally thought to possess it (*2, 5*). This opens up the possibility that the study of the diverse forms of mammalian vocal learning behaviors could broaden our understanding of the core molecular, anatomical and physiological brain mechanisms of vocal learning as well as the mechanisms underlying the convergent evolution of skilled motor behaviors more broadly.

**Figure 1.**
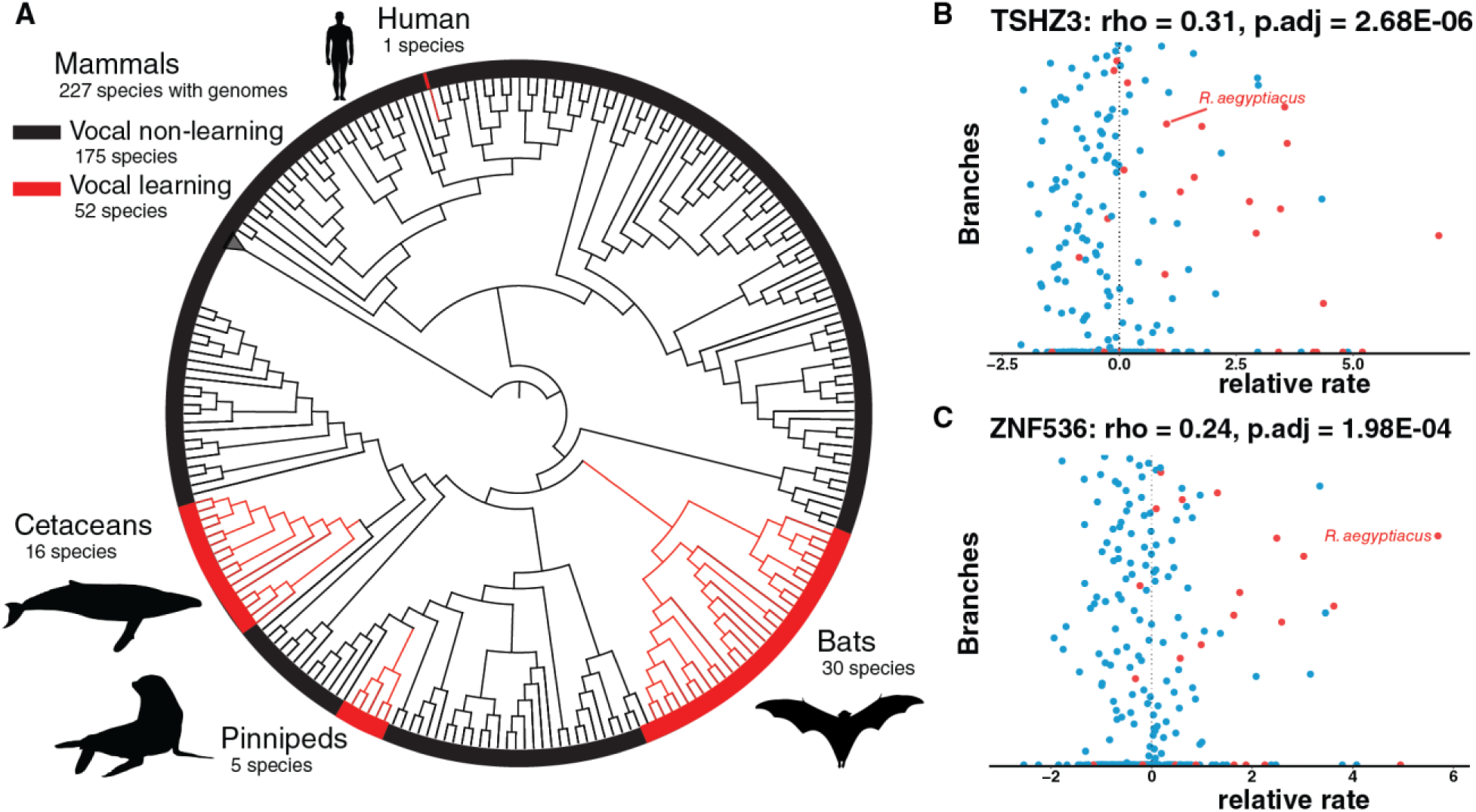
**(A)** A cladogram of mammalian species whose genomes were analyzed in this study highlights the convergent evolution of vocal learning species (in red) relative to non-learners (in black). The phylogenetic tree used in our analyses was derived from a companion paper to this study (*63*). Both *TSHZ3* **(B)** and *ZNF536* **(C)** exhibit relative evolutionary rate (RER) shifts in vocal learning mammals (red) relative to vocal non-learners (blue).

We sought to evaluate the presence of convergent genomic specializations between vocal learning mammals using new datasets and computational approaches, focusing on bats as an attractive mammalian model of vocal complexity (*6–13*). Specifically, we used protein-coding sequences from genomes generated by the Zoonomia Consortium (*14, 15*) and models of evolutionary rate convergence (*16*)to identify genes repeatedly associated with the evolution of vocal learning. Motivated by the finding of protein-level convergence, we next profiled open chromatin specializations of multiple brain regions and somatic tissues in the Egyptian fruit bat, a bat with robust vocal plasticity (*10–12*), to identify vocalization-associated epigenomic specializations. We were able to accomplish this by combining anatomical tracing and electrophysiological recordings in vocalizing bats to identify a region of motor cortex associated with vocal production. The vocalization-associated epigenomic data collected from bat– combined with the availability of hundreds of mammalian genomes (*17*), their associated reference-free whole-genome alignments (*18*), and high-quality epigenomic data from motor cortex of multiple additional mammalian species (*19–21*)–provided the foundation for us to apply a new machine learning approach developed in our companion paper, the Tissue-Aware Conservation Inference Toolkit (TACIT) (*22*), to identify putative enhancer sequences associated with the convergent evolution of vocal learning. In sum, we find evidence of convergent genetic evolution across mammals in both protein coding and non-coding DNA sequences by leveraging the diversity in mammal vocal learning behavior, novel computational tools we developed, and new anatomical, electrophysiological, and regulatory genomics measurements for a bat with robust vocal plasticity.

### Convergent Evolution in Protein Sequence Associated with Vocal Learning Behavior

To explore the possibility of shared genomic specializations associated with vocal learning, we first used new protein-coding alignments for hundreds of mammals (*14*) to identify genes whose rates of evolution differs between vocal learners and other mammals, and which may thus be under evolutionary selection related to vocal learning (*16*). We analyzed 16,209 high-quality gene alignments across 175 boreoeutherian mammals, including 25 vocal learning species (Materials and Methods). We found evidence for differential evolutionary rates between vocal learners and non-learners in 865 genes (p.adj < 0.01), with a smaller set of 201 genes being robust to dropout of each of the vocal learning lineages (Data S1). Gene ontology analysis revealed terms associated with transcription as the topmost enriched functional pathways in these convergent gene sets, supporting gene regulation as a critical process in the evolution of mammalian vocal learning (Data S2). Notably, two of these genes—*TSHZ3* and *ZNF536* (Fig. 1B,C)—are syntenic neighboring loci on human chromosomal region 19q12, which in humans has been strongly linked to severe speech disability (*23–25*).

### Identification of a Vocal Production Region in Egyptian Fruit Bat

The enrichment of transcription factors in the set of vocal learning-associated proteins suggests that differences in gene regulation are likely to be driving the evolution of vocal learning. Since gene regulation is often tissue-specific, we next sought to identify regions of the brain involved in vocal production. For this, we compared brain regions involved in speech production in humans to corresponding regions in the Egyptian fruit bat, *Rousettus aegyptiacus*, a bat species with robust vocal plasticity (*12, 26*). To identify a candidate region, we were guided by the notion that fine vocal-motor control may be associated with anatomical specialization of the motor cortex (*27–31*). In particular, previous work suggested that a cortical region controlling complex vocal behavior would be characterized by a direct, monosynaptic projection onto the motoneurons controlling the vocal source (in mammals, the larynx). Such a direct connection has been observed robustly in vocal learning birds (songbirds, parrots and hummingbirds, (*32–34*)) and suggested in mice (*35*), chimpanzees (*36*), and humans (*37–40*).

Therefore, to guide our anatomical identification of a vocal area in the bat motor cortex, we first determined whether a direct corticospinal anatomical connection exists in *R. aegyptiacus*. Guided by cortical mapping experiments (*41*), we injected anterograde tracers into part of the motor cortex that has been associated with orofacial motor control (ofM1) and searched for labeled descending cortical fibers in the hindbrain region where the laryngeal motoneurons reside, the nucleus ambiguus (NA) (Fig. 2A, Supp. Fig. 1A-B and Supp. Movie 1). To test the existence of a direct monosynaptic projection, we also specifically identified laryngeal motoneurons in the NA by retrogradely labeling them through bilateral muscular injection of CTB into the cricothyroid muscles of the bat larynx (Fig. 2A). We validated the colocalization of descending cortical fibers and local synaptic boutons with laryngeal motoneurons using two complementary labeling approaches, one relying on immunostaining of synapses (VGLUT1) and the other one using viral labeling of synapses (SYN) (Fig. 2B-F; Supp. Fig. 2). Across five bats, 79.2% of the retrogradely labeled motoneurons (61/77) colocalized with descending cortical fibers and 26% of them (20/77) colocalized with both cortical fibers and synaptic boutons, pointing to the existence of a robust direct corticospinal projection to laryngeal motoneurons (Fig. 2G). This colocalization in the NA was not only consistent across the different techniques (Fig. 2G, Supp. Fig. 2) but could not be found in any other brainstem motor nuclei, including the Hypoglossal nucleus, which controls the tongue and neck muscles (Supp. Fig. 1C-E). Interestingly, the cortico-bulbar fibers crossed the midline anterior to the NA at the level of the facial nucleus, offering a direct contra-lateral path for the innervation of the NA (Supp. Fig. 1F). These anatomical findings highlight the bat ofM1 as a possible candidate region associated with vocal production.

**Figure 2.**
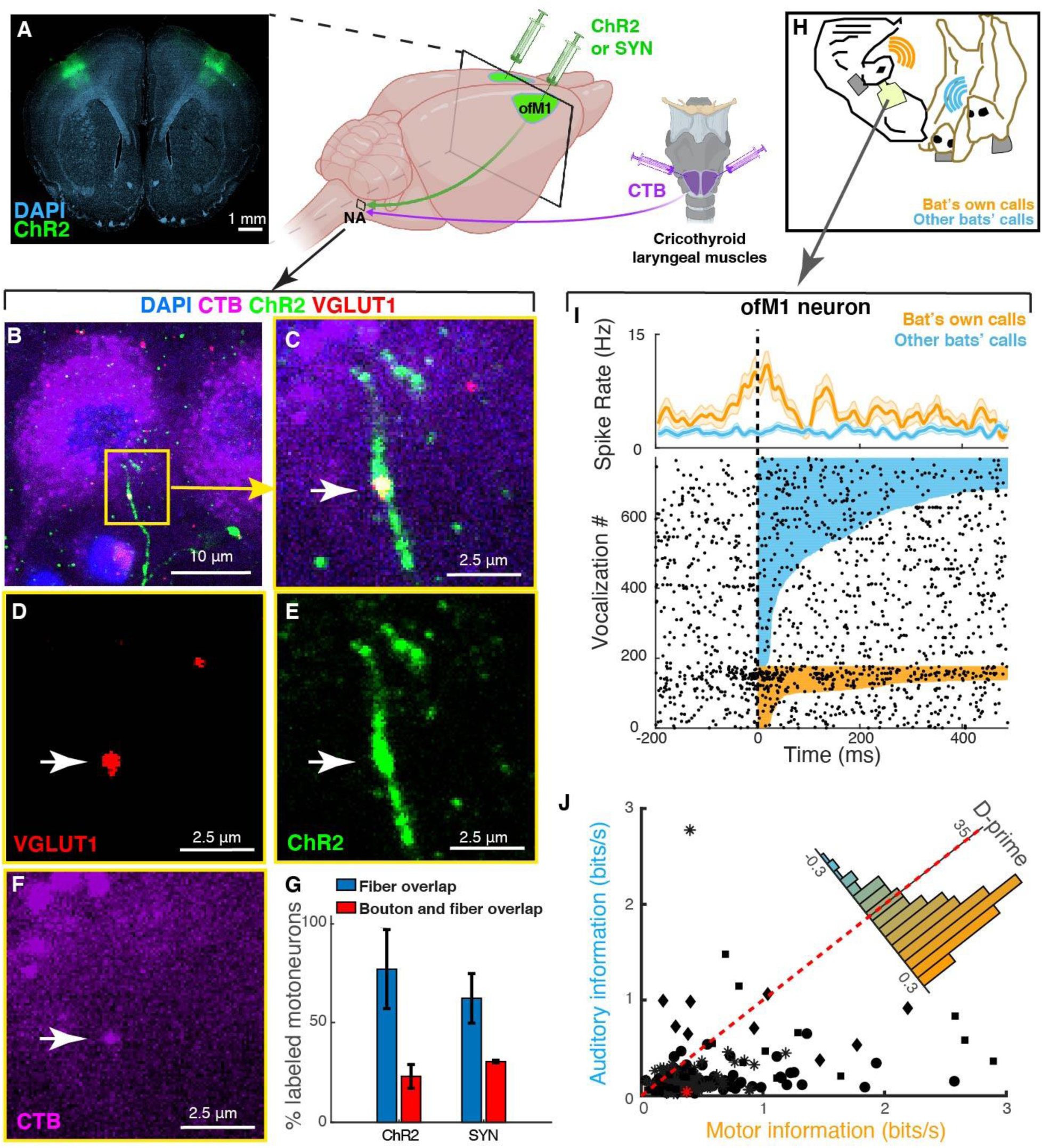
Identification of an anatomically specialized motorcotical region targeting laryngeal motoneurons in the Egyptian fruit bat. **(A)** Right: schematic of anatomical tracing approaches. Retrograde tracer cholera toxin B (CTB, purple) was injected bilaterally into the cricothyroid muscles to label brainstem motoneurons in nucleus ambiguus (NA). Simultaneously, an anterograde tracer (channelrhodopsin-2, ChR2, or Synpasin, SYN; green) was injected bilaterally into the orofacial motor cortex (ofM1) to label corticobulbar projections into NA. Left: example coronal section showing cortical injection sites with anterograde tracer (ChR2, green) and DAPI labeling (cyan). **(B-F)** Laryngeal motoneurons in the NA identified using a retrograde tracer (CTB, purple), cortical fibers labeled with ChR2 (green), corticobulbar synapses labeled with VGLUT1 (red), and DAPI (blue). **B** and **C** are overlaid images showing colocalization of fibers with a synaptic bouton on the retrograde labeled cell (white arrow). **(G)** Percentage of laryngeal motoneurons labeled with CTB that are colocalized with cortical fibers (blue) or with both cortical fibers and synaptic boutons (red). Note that both tracing techniques qualitatively yielded similar results: ChR2, *n* = 51 cells from 3 bats; Synapsin/synaptophysin dual-label virus (SYN), *n* = 26 cells from 2 bats. **(H)** Illustration of the experimental setup during which wireless electrophysiological recordings were conducted from the identified cortical region in freely behaving and vocalizating bats. **(I)** Spiking activity of an example ofM1 neuron aligned to the onset of vocalizations produced (bat’s own calls, orange) or heard (other bats’ calls, blue) by the bat subject. *Top row*, time varying mean firing rate and corresponding raster plot below. Colored lines in the raster plot show the duration of each vocalization. Note increased firing rate during vocal production as compared to hearing. **(J)** Information (see Methods) between the time varying firing rate and the amplitude of produced (x-axis) vs. heard (y-axis) vocalizations for 219 single units (marker shapes indicate bat ID, *n*=4 bats). The cell shown in **(I)** is highlighted in red. Inset shows the distribution of D-prime between motor and auditory information for the same cells. Note that the distribution is heavily skewed towards higher motor information rather than auditory information coded in the activity of the recorded neurons. Error bars are mean +/- SEM throughout the figure.

To further corroborate the role of ofM1 in vocal control, we tested whether ongoing neural activity in this area was associated with vocal production. As single unit recordings have never been performed from an identified laryngeal region of the motor cortex in any mammalian species, including in humans, we tested whether this region indeed exhibited modulation of neural activity during vocal behavior. We performed wireless electrophysiological recordings from four bats engaged in free vocal interactions with peers (Fig. 2H). Vocalizations were identified and recorded using wireless call detectors placed around the necks of the individual bats (see Materials and Methods, (*26*)). We found that about half of the recorded single units in ofM1 (115/237) showed a significant change in firing rates when the bats produced vocalizations as compared to staying quiet (Supp. Fig. 3A-C; Anova on Poisson GLM per cell; *p-value* threshold = 0.001; Materials and Methods). Importantly, in 24.5% of ofM1 cells that were excited during vocal production (26/106), the change of activity could not be accounted for by jaw or tongue movements (Supp. Fig. 3D). Furthermore, many of the single units had a sustained increase of activity during production but not during perception of vocalizations (Fig. 2I). To further assess this specific neural modulation during vocal-motor production, we quantified the information between the time varying firing rate and the amplitude modulation of the vocalizations. This analysis confirmed that ofM1 neurons had significantly higher motor than auditory information (Fig. 2J; likelihood-ratio test on LME models, N=219, LRStat = 62.515, df = 1, p = 2.6645x10^-5^; average d-prime change in information gain during motor production = 0.15 ± 0.13, corresponding to an increase of 0.286 ± 0.035 bits/s). Combined, the results of the anatomical and electrophysiological study defined ofM1 as a motor cortical area associated with vocal behavior in *R. aegyptiacus*.

**Figure 3.**
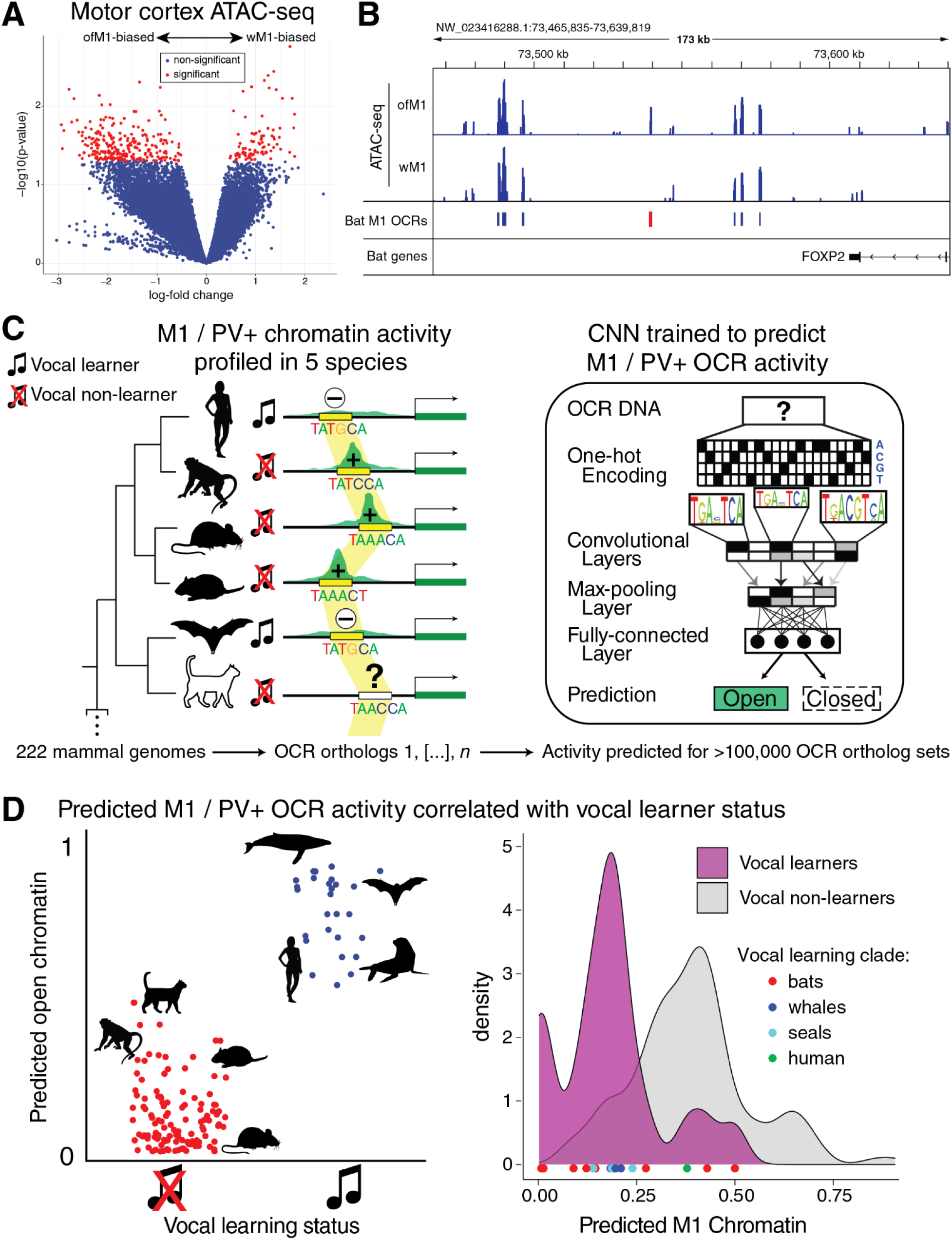
A machine learning approach enables the identification of open chromatin regions (OCRs) associated with vocal learning in mammals. **(A)** Volcano plot of ATAC-seq OCRs with differential activity between the orofacial and wing subregions of primary motor cortex (ofM1 and wM1, respectively) of Egyptian fruit bat. **(B)** Genomic browser showing ofM1 and wM1 ATAC-seq traces at the 3’ end of the FOXP2 locus. Reproducible M1 open chromatin regions (OCRs) are indicated in blue, with a differentially active OCR in ofM1 relative to wM1 highlighted in red. **(C)** OCRs identified in motor cortex (M1, 4 species) or M1 parvalbumin-positive neurons (PV, 2 species) were mapped across 222 mammalian genomes (left) to train convolutional neural networks to predict M1 or PV open chromatin from sequence alone. **(D)** Using the Tissue-Aware Conservation Inference Toolkit (TACIT (*22*)), OCRs were identified whose predicted open chromatin status across species was significantly associated with those species’ vocal learning status (left), as in the case of an OCR proximal to *TSHZ3* and *ZNF536* (right).

### Epigenomic Specializations in the Vocal Production Region of the Egyptian Fruit Bat Motor Cortex

We next sought to epigenomically profile vocal and non-vocal brain regions in *R. aegyptiacus* in order to identify vocal learning-associated regulatory genomic specializations. We first generated a multi-tissue atlas of open chromatin data—indicative of regulatory activity—by performing ATAC-seq (assay for transposase-accessible chromatin sequencing (*42*)) across 7 brain regions and 3 somatic tissues and of *R. aegyptiacus* (Materials and Methods), including ofM1. From a total set of 88,389 non-coding, non-promoter open chromatin regions (OCRs) in primary motor cortex (M1), we identified 348 candidate enhancers with differential open chromatin between orofacial motor cortex (ofM1) and wing motor cortex (wM1) (Fig. 3A, Data S3, Materials and Methods). Genes proximal to OCRs with differential epigenomic activity between ofM1 and wM1 were significantly enriched for functional association with neuronal projections and transcriptional regulation (Data S4). These included OCRs near 51 known transcription factors (TFs), including *FOXP2*, a TF that has been extensively implicated in human speech and vocal learning (Fig. 3B) (*43*). Notably, genes near OCRs differentially open between bat ofM1 and wM1 included genes we had also identified as being under convergent acceleration in vocal learners (n = 5) (Data S1). These specialized regions of open chromatin, coupled with enrichment of transcription factors in the set of vocal learning-associated protein-coding genes, both suggest that both *cis* and *trans* differences in gene regulation are driving forces in the evolution of vocal learning behaviors.

### Convergent Evolution in Candidate Enhancer Sequences Associated with Vocal Learning Behavior

Since there is increasing evidence that *cis*-regulatory differences in enhancer regions are driving the evolution of complex traits (*44–46*), we sought to identify OCRs associated with the evolution of vocal learning. Detecting *cis*-regulatory element differences associated with trait evolution is challenging because many regulatory enhancers can preserve the same regulatory function even when the underlying genome sequence is highly divergent and many *cis*-regulatory elements have tissue-specific activity (*47–49*) Thus, methods for convergent evolution that rely on the alignment of individual nucleotides between species (e.g. (*16, 50, 51*)) are likely to miss a substantial proportion of key candidate enhancers.

We therefore sought to extend our search for cis-regulatory elements whose evolution is associated with vocal learning behavior using a new machine learning approach, TACIT (Tissue-Aware Conservation Inference Toolkit (*22*)). Given that it is infeasible to map the brains and collect motor cortex tissue from each vocal learning and closely related non-learning species, the TACIT approach uses machine learning models (*52*) to predict motor cortex open chromatin, across orthologous regions of the genome (*47–49*). TACIT then associates predictions with vocal learning in a way that corrects for phylogenetic relationships (Fig. 3C). We leveraged DNA sequence-based M1 open chromatin predictions from convolutional neural networks (CNNs) trained on our ATAC-seq data from *R. aegyptiacus* M1 and previously collected data from M1 of mouse (*20*), rat, and macaque (*19*) to predict motor cortex open chromatin across 222 mammalian genomes (Materials and Methods, (*22*)). Given that Parvalbumin has been shown to be a shared marker of brain areas critical for vocal learning in song-learning birds and humans (*4*), we also used CNNs trained to predict cell type-specific OCR activity using ATAC-seq data from mouse and human M1 Parvalbumin-positive neurons (M1-PV+) (*21, 22, 53*).

We subsequently used phylogenetic logistic regression (*54, 55*) with phylogenetic permulations (*56*) to identify OCRs whose predicted open chromatin differences across species showed convergent similarities in vocal learning mammals relative to non-learners (Fig. 3D, Materials and Methods, (*22*)). We identified 38 candidate enhancers from our M1 CNN models (Table S1) and 10 from our M1-PV+ neuron models (Table S2) that displayed convergent patterns of predicted open chromatin in vocal learning species. In the majority of cases, the genes closest to these putative enhancers have been associated with significant developmental delay or complete absence of speech when disrupted in humans (Tables S1-2). Five of the OCRs identified by the M1 model were proximal to genes—*ERBB4*, *GALC*, *TCF4*, *TSHZ3*, and *ZNF536*—that were also near OCRs with differential activity between bat ofM1 and wM1, as well as three genes with convergent evolutionary acceleration in vocal learning mammals (Fig. 4, Data S1). Two of the vocal learning-associated M1 OCRs were proximal to genes—*DAAM1* and *VIP*—which were previously shown to have convergent gene regulation between humans and song-learning birds (*4*). Consistent with the finding that convergent vocal learning-associated gene regulation is primarily repressive (*4*), we found the majority of candidate enhancers (n = 29/38 OCRs, 76%) had lower predicted open chromatin activity in the vocal learning mammals relative to controls.

**Figure 4.**
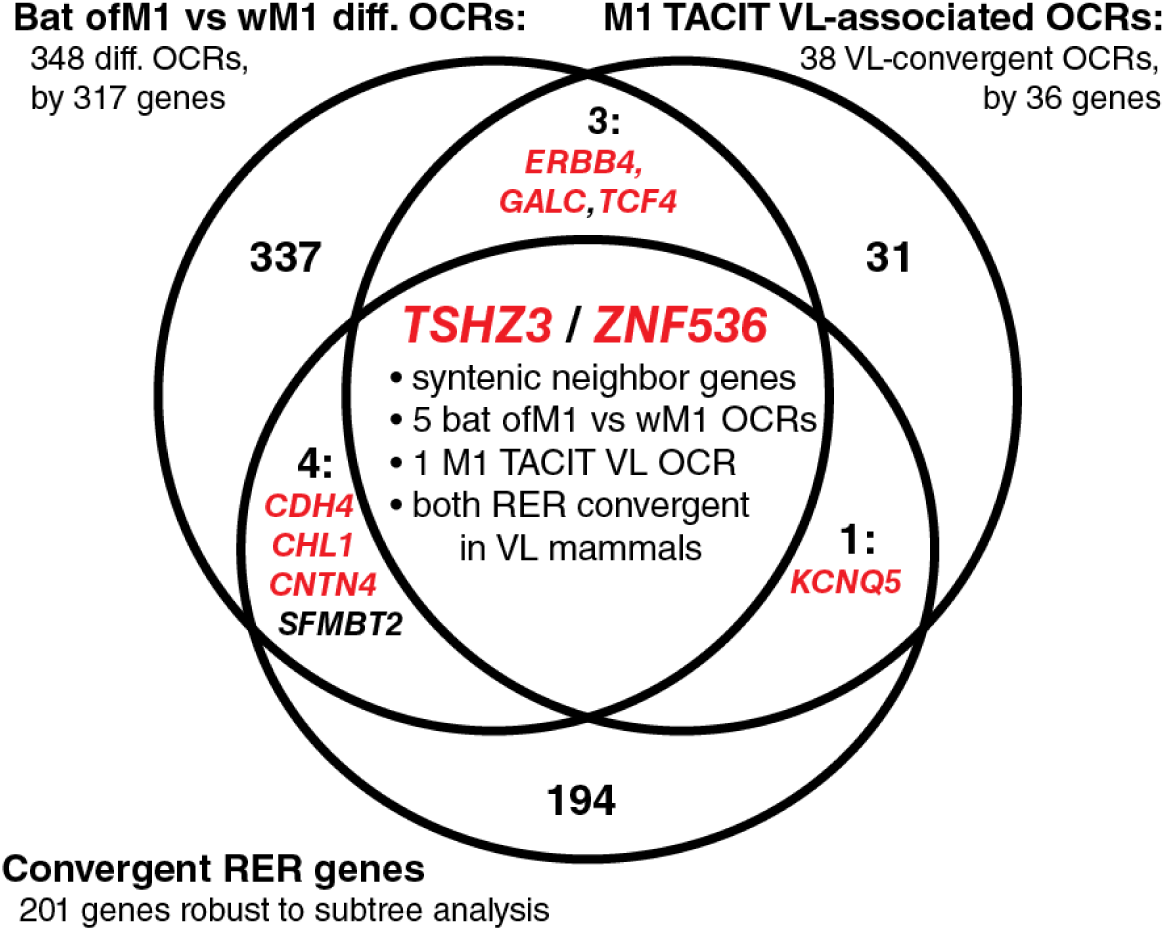
Multiple lines of evidence support *TSHZ3* and *ZNF536* as top candidate genes associated with vocal learning in mammals. Overlap between sets of genes proximal to differential open chromatin regions (OCRs) between orofacial (ofM1) and wing (wM1) motor cortices in a bat, genes proximal to OCRs predicted by TACIT to have convergent M1 open chromatin in vocal learning mammals, and genes whose protein-coding sequences are under RER convergence in vocal learning mammals. Genes whose disruption has been associated with speech disability in humans are indicated in red (references in Table S1, except for *CDH4* (*64*), *CHL1* (*65*), and *CNTN4* (*66*)).

To explore the possibility that higher-level gene regulatory networks could be driving specialized gene expression related to vocal motor behavior, we identified transcription factor motifs enriched in OCRs that displayed differential activity between bat ofM1 and wM1 (Data S5). The most enriched TF motif in the set of ofM1-specialized OCRs was that of *ETS1*, a member of a TF family that has been implicated in the formation of selective projections between motor neurons and their muscle targets (*57*). Notably, the *ETS1* locus was itself also associated with an OCR with differential activity between ofM1 and wM1. The second-most enriched motif was that of *NFATC2*, a TF that has been shown to form a co-binding complex with speech-associated gene *FOXP2* (*58*). Incidentally, the *NFATC2* locus was also associated with an M1 OCR predicted by TACIT to be convergently regulated in vocal learning mammals. The finding of TF motifs enriched in OCRs that themselves may be regulating downstream TFs suggests a role for complex higher-order gene regulatory mechanisms in the evolution of vocal learning.

Multiple lines of evidence suggest a model in which differences in gene regulatory networks of transcription factors mediate behavioral differences in vocal learning across clades. First, we found evidence of vocal learning-associated convergent evolution in 201 protein coding genes, which are enriched for gene regulatory functions. Second, differential regions of open chromatin between the orofacial and wing subregions of the bat motor cortex implicate networks of transcription factors as critical *cis*-regulatory targets in this region. Third, TACIT identified 48 candidate enhancers whose predicted open chromatin state across species is associated with vocal learning behavior. Those differences are likely due to gains and losses of transcription factor binding sites at those regions of open chromatin (*22*). These findings are consistent with previous work that FOXP2 (*43*) and other transcription factors like NEUROD6 and the MEF2 family (*59*) are involved in vocal learning ability in humans and songbirds. Although our *cis*-regulatory results clearly implicate motor cortex specializations, we cannot rule out that these candidate enhancers and their associated vocal learning-associated genes impact a number of different brain regions and cell types.

Our results provide a foundation for studying the convergent evolution of molecular mechanisms for vocal learning. Despite relying on different input data and orthogonal methodologies, we found agreement in the results between the different approaches we took to identify vocal learning-associated loci, in particular at the locus containing neurodevelopmental transcription factors TSHZ3 and ZNF536 (Fig. 4). Although these genes have not previously been appreciated for their role in vocal learning, the 19q12 region of the human genome that contains them has been repeatedly implicated in severe speech disability (*23–25*). We found the bat ortholog of this locus to contain 5 OCRs—more than any other locus—with enhanced activity in ofM1 relative to wM1, as well as a candidate enhancer significantly associated with vocal learning across mammals by our M1 TACIT model. Further, we found that both the *TSHZ3* and *ZNF536* genes themselves were under convergent evolutionary acceleration in vocal learning mammals. In humans, disruption of *TSHZ3* impacts respiratory rhythms and cortico-striatal circuits, which are both involved in the learning and production of vocalizations (*60, 61*). ZNF536 has been implicated in the transcriptional inhibition of genes involved in neuronal differentiation, with depletion or mutation resulting in an enhancement of neuronal differentiation (*62*).

Vocal learning has convergently evolved in multiple lineages of mammals, allowing us to leverage new mammalian genomes to identify shared genomic specializations associated with the behavior. We find evidence at multiple levels that the differences in vocal learning in behavior across species are associated with convergent differences in genome sequence, largely centered around higher-level transcriptional regulation. However, the extent to which these sequence differences are causal, rather than a consequence, of behavioral evolution remains an open question. Overall, our results provide a foundation for studying the genomic underpinnings of vocal learning in mammals and open up exciting future applications, including the extension of our models to birds and the possibility of predicting whether unclassified species are vocal learners. Our novel approach of integrating amino acid, neural circuit, and *cis*-regulatory evolution using machine learning has the potential to be applied to a variety of different convergently evolved traits and behaviors.

## Supporting information

Data_S1_VL_RERconverge_gene_sets

Data_S2_VL_RERconverge_gene_functional_enrichments

Data_S3_ofM1_wM1_differential_OCRs

Data_S4_OCR_gene_functional_enrichments

Data_S5_OCR_TF_motif_enrichments

Data_S6_VL_species_annotations

## Acknowledgments

We would like to thank Andrew C. Halley and Leah Krubitzer for helpful discussions and critical assistance in identifying orofacial and wing subregions of the *Rousettus aegyptiacus* motor cortex. We would like to thank Frederic E. Theunissen for helpful discussions around the analysis of coherence of neural activity and for sharing his lab’s code. We would like to thank the other members of the Zoonomia Consortium, the Vertebrate Genomes Project Vocal Learning working group, and the Pfenning and Yartsev labs for useful discussions and suggestions. Figure 2A was created with BioRender.com.

## Funding

Alfred P. Sloan Foundation Research Fellowship (ARP)

National Institutes of Health grant NIDA DP1DA046585 (ARP)

National Institutes of Health grant R01NS111479 (EA)

National Institutes of Health grant R01 NS121231 (BL)

National Science Foundation grant NSF IOS-2022241 (BL).

National Institutes of Health grant DP2 DP2-DC016164 (MMY).

The New York Stem Cell Foundation NYSCF-R-NI40 (MMY).

Alfred Sloan Foundation FG-2017-9646 (MMY).

Brain Research Foundation BRFSG-2017-09 (MMY).

Packard Foundation Fellowship 2017-66825 (MMY).

Klingestein-Simons Fellowship (MMY).

Human Frontiers Research Program (MMY).

Pew Charitable trust 00029645 (MMY).

McKnight Foundation 042823 (MMY).

Dana Foundation (MMY).This work used the Extreme Science and Engineering Discovery Environment (XSEDE), through the Pittsburgh Supercomputing Center Bridges and Bridges-2 Compute Clusters, which was supported by the National Science Foundation (grant TG-BIO200055).

## Author Contributions

Conceptualization: MEW, ARP, TAS, JEE, MMY

Methodology: MEW, ARP, TAS, JEE, MMY, IMK

Resources: TAS, MMY, WM, EA, BKL

Data Curation: JEE, XZ

Investigation: MEW, TAS, JEE, VAS, AR, MBJ, NSB, MMY

Software: MEW, XZ, IMK, DES, SA, JEE

Formal analysis: MEW, XZ, IMK, DES, AJL, ARP, MMY, TAS, JEE, VAS, AR, MBJ, NSB

Funding acquisition: ARP, MMY

Supervision: ARP, MEW, MMY, TAS, IMK

Writing – original draft: MEW, TAS, JEE, ARP

Writing – review & editing: MEW, TAS, JEE, IMK, DES, AJL, MMY, ARP

## Competing interests

Authors declare that they have no competing interests.

## Data and materials availability

Egyptian fruit bat ATAC-seq data—including raw .fasta files, .bigWig files of genome-aligned reads, processed open chromatin .narrowpeak files, and extensive sample metadata—have been made available at GEO under accession ID GSE187366. The histological and electrophysiology data set from this study are available from the corresponding authors upon request.

## Zoonomia Consortium Members

Gregory Andrews^1^; Joel C. Armstrong^2^; Matteo Bianchi^3^; Bruce W. Birren^4^; Kevin R. Bredemeyer^5^; Ana M. Breit^6^; Matthew J. Christmas^3^; Hiram Clawson^2^; Joana Damas^7^; Federica Di Palma^8,9^; Mark Diekhans^2^; Michael X. Dong^3^; Eduardo Eizirik^10^; Kaili Fan^1^; Cornelia Fanter^11^; Nicole M. Foley^5^; Karin Forsberg-Nilsson^12,13^; Carlos J. Garcia^14^; John Gatesy^15^; Steven Gazal^16^; Diane P. Genereux^4^; Daniel Goodman^17^; Linda Goodman^18^; Jenna Grimshaw^14^; Michaela K. Halsey^14^; Andrew J. Harris^5^; Glenn Hickey^19^; Michael Hiller^20,21,22^; Allyson G. Hindle^11^; Robert M. Hubley^23^; Laura M. Huckins^24^; Graham M. Hughes^25^; Jeremy Johnson^4^; David Juan^26^; Irene M. Kaplow^27,28^; Elinor K. Karlsson^1,4,29^; Kathleen C. Keough^18,30,31^; Bogdan Kirilenko^20,21,22^; Klaus-Peter Koepfli^32,33,34^; Jennifer M. Korstian^14^; Amanda Kowalczyk^27,28^; Sergey V. Kozyrev^3^; Alyssa J. Lawler^4,28,35^; Colleen Lawless^25^; Danielle L. Levesque^6^; Harris A. Lewin^7,36,37^; Xue Li^1,4,38^; Yun Li^39^; Abigail Lind^30,31^; Kerstin Lindblad-Toh^3,4^; Ava Mackay-Smit^40^; Voichita D. Marinescu^3^; Tomas Marques-Bonet^41,42,43,44^; Victor C. Mason^45^; Jennifer R. S. Meadows^3^; Wynn K. Meyer^46^; Jill E. Moore^1^; Lucas R. Moreira^1,4^; Diana D. Moreno-Santillan^14^; Kathleen M. Morrill^1,4,38^; Gerard Muntané^26^; William J. Murphy^5^; Arcadi Navarro^41,43,47,48^; Martin Nweeia^49,50,51,52^; Austin Osmanski^14^; Benedict Paten^2^; Nicole S. Paulat^14^; Eric Pederson^3^; Andreas R. Pfenning^27,28^; BaDoi N. Phan^27,28,53^; Katherine S. Pollard^30,31,54^; Kavya Prasad^27^; Henry Pratt^1^; David A. Ray^14^; Steven K. Reilly^40^; Jeb R. Rosen^23^; Irina Ruf^55^; Louise Ryan^25^; Oliver A. Ryder^56,57^; Daniel E. Schäffer^27^; Aitor Serres^26^; Beth Shapiro^58,59^; Arian F. A. Smit^23^; Mark Springer^60^; Chaitanya Srinivasan^27^; Cynthia Steiner^56^; Jessica M. Storer^23^; Kevin A. M. Sullivan^14^; Patrick F. Sullivan^39,61^; Quan Sun^62^; Elisabeth Sundström^3^; Megan A. Supple^58^; Ross Swofford^4^; Jin Szatkiewicz^39^; Joy-El Talbot^63^; Emma Teeling^25^; Jason Turner-Maier^4^; Alejandro Valenzuela^26^; Franziska Wagner^64^; Ola Wallerman^3^; Chao Wang^3^; Juehan Wang^16^; Jia Wen^39^; Zhiping Weng^1^; Aryn P. Wilder^56^; Morgan E. Wirthlin^27,28,65^; James R. Xue^4,66^; Shuyang Yao^61^; Xiaomeng Zhang^4,27,28^

## Affiliations

^1^Program in Bioinformatics and Integrative Biology, UMass Chan Medical School; Worcester, MA 01605, USA

^2^Genomics Institute, University of California Santa Cruz; Santa Cruz, CA 95064, USA

^3^Department of Medical Biochemistry and Microbiology, Science for Life Laboratory, Uppsala University; Uppsala, 751 32, Sweden

^4^Broad Institute of MIT and Harvard; Cambridge, MA 02139, USA

^5^Veterinary Integrative Biosciences, Texas A&M University; College Station, TX 77843, USA

^6^School of Biology and Ecology, University of Maine; Orono, ME 04469, USA

^7^The Genome Center, University of California Davis; Davis, CA 95616, USA

^8^Genome British Columbia; Vancouver, BC, Canada

^9^School of Biological Sciences, University of East Anglia; Norwich, UK

^10^School of Health and Life Sciences, Pontifical Catholic University of Rio Grande do Sul; Porto Alegre, 90619-900, Brazil

^11^School of Life Sciences, University of Nevada Las Vegas; Las Vegas, NV 89154, USA

^12^Biodiscovery Institute, University of Nottingham; Nottingham, UK

^13^Department of Immunology, Genetics and Pathology, Science for Life Laboratory, Uppsala University; Uppsala, 751 85, Sweden

^14^Department of Biological Sciences, Texas Tech University; Lubbock, TX 79409, USA

^15^Division of Vertebrate Zoology, American Museum of Natural History; New York, NY 10024, USA

^16^Keck School of Medicine, University of Southern California; Los Angeles, CA 90033, USA

^17^Department of Immunology, University of California San Francisco; San Francisco, CA 94143, USA

^18^Fauna Bio Incorporated; Emeryville, CA 94608, USA

^19^Baskin School of Engineering, University of California Santa Cruz; Santa Cruz, CA 95064, USA

^20^Faculty of Biosciences, Goethe-University; 60438 Frankfurt, Germany

^21^LOEWE Centre for Translational Biodiversity Genomics; 60325 Frankfurt, Germany

^22^Senckenberg Research Institute; 60325 Frankfurt, Germany

^23^Institute for Systems Biology; Seattle, WA 98109, USA

^24^Department of Psychiatry, Icahn School of Medicine at Mount Sinai; New York, NY 10029, USA

^25^School of Biology and Environmental Science, University College Dublin; Belfield, Dublin 4, Ireland

^26^Department of Experimental and Health Sciences, Institute of Evolutionary Biology (UPF-CSIC), Universitat Pompeu Fabra; 08003, Barcelona, Spain

^27^Department of Computational Biology, School of Computer Science, Carnegie Mellon University; Pittsburgh, PA 15213, USA

^28^Neuroscience Institute, Carnegie Mellon University; Pittsburgh, PA 15213, USA

^29^Program in Molecular Medicine, UMass Chan Medical School; Worcester, MA 01605, USA

^30^Department of Epidemiology & Biostatistics, University of California San Francisco; San Francisco, CA 94158, USA

^31^Gladstone Institutes; San Francisco, CA 94158, USA

^32^Center for Species Survival, Smithsonian Conservation Biology Institute, National Zoological Park; Washington, DC 20008, USA

^33^Computer Technologies Laboratory, ITMO University; St. Petersburg 197101, Russia

^34^Smithsonian-Mason School of Conservation; Front Royal, VA 22630, USA

^35^Department of Biological Sciences, Mellon College of Science, Carnegie Mellon University; Pittsburgh, PA 15213, USA

^36^Department of Evolution and Ecology, University of California Davis; Davis, CA 95616, USA

^37^John Muir Institute for the Environment, University of California Davis; Davis, CA 95616, USA

^38^Morningside Graduate School of Biomedical Sciences, UMass Chan Medical School; Worcester, MA 01605, USA

^39^Department of Genetics, University of North Carolina Medical School; Chapel Hill, NC 27599, USA

^40^Department of Genetics, Yale School of Medicine; New Haven, CT 06510, USA

^41^Catalan Institution of Research and Advanced Studies (ICREA); 08010, Barcelona, Spain

^42^CNAG-CRG, Centre for Genomic Regulation, Barcelona Institute of Science and Technology (BIST); 08036, Barcelona, Spain

^43^Department of Medicine and LIfe Sciences, Institute of Evolutionary Biology (UPF-CSIC), Universitat Pompeu Fabra; 08003, Barcelona, Spain

^44^Institut Català de Paleontologia Miquel Crusafont, Universitat Autònoma de Barcelona; 08193, Cerdanyola del Vallès, Barcelona, Spain

^45^Institute of Cell Biology, University of Bern; 3012, Bern, Switzerland

^46^Department of Biological Sciences, Lehigh University; Bethlehem, PA 18015, USA

^47^BarcelonaBeta Brain Research Center, Pasqual Maragall Foundation; Barcelona, 08005, Spain

^48^CRG, Centre for Genomic Regulation, Barcelona Institute of Science and Technology (BIST); 08003, Barcelona, Spain

^49^Department of Comprehensive Care, School of Dental Medicine, Case Western Reserve University; Cleveland, OH 44106, USA

^50^Department of Vertebrate Zoology, Canadian Museum of Nature; Ottawa, Ontario K2P 2R1, Canada

^51^Department of Vertebrate Zoology, Smithsonian Institution; Washington, DC 20002, USA

^52^Narwhal Genome Initiative, Department of Restorative Dentistry and Biomaterials Sciences, Harvard School of Dental Medicine; Boston, MA 02115, USA

^53^Medical Scientist Training Program, University of Pittsburgh School of Medicine; Pittsburgh, PA 15261, USA

^54^Chan Zuckerberg Biohub; San Francisco, CA 94158, USA

^55^Division of Messel Research and Mammalogy, Senckenberg Research Institute and Natural History Museum Frankfurt; 60325 Frankfurt am Main, Germany

^56^Conservation Genetics, San Diego Zoo Wildlife Alliance; Escondido, CA 92027, USA

^57^Department of Evolution, Behavior and Ecology, School of Biological Sciences, University of California San Diego; La Jolla, CA 92039, USA

^58^Department of Ecology and Evolutionary Biology, University of California Santa Cruz; Santa Cruz, CA 95064, USA

^59^Howard Hughes Medical Institute, University of California Santa Cruz; Santa Cruz, CA 95064, USA

^60^Department of Evolution, Ecology and Organismal Biology, University of California Riverside; Riverside, CA 92521, USA

^61^Department of Medical Epidemiology and Biostatistics, Karolinska Institutet; Stockholm, Sweden

^62^Department of Biostatistics, University of North Carolina at Chapel Hill; Chapel Hill, NC, USA

^63^Iris Data Solutions, LLC; Orono, ME 04473, USA

^64^Museum of Zoology, Senckenberg Natural History Collections Dresden; 01109 Dresden, Germany

^65^Allen Institute for Brain Science; Seattle, WA 98109, USA

^66^Department of Organismic and Evolutionary Biology, Harvard University; Cambridge, MA 02138, USA

## Supplementary Materials

### Materials and Methods

Supplementary Text

Tables S1 to S2

Data S1 to S6

Figures S1 to S3

Movie S1

References (67–129)

## Supplementary Materials

This PDF file includes:

## Materials and Methods

Tables S1 to S2

Figures S1 to S3

### Other Supplementary Materials for this manuscript include the following

Data_S1_VL_RERconverge_gene_sets.xlsx Data_S2_VL_RERconverge_gene_functional_enrichments.xlsx

Data_S3_ofM1_wM1_differential_OCRs.xlsx

Data_S4_OCR_gene_functional_enrichments.xlsx

Data_S5_OCR_TF_motif_enrichments.xlsx

Data_S6_VL_species_annotations.xlsx

Movie 1.mp4

## Materials and Methods

### Coding the vocal learning trait

We annotated the vocal learning trait for a set of 215 mammalian species that satisfied the following conditions: (1) the species’ genome was present in the Zoonomia whole-genome Cactus alignment (241 assemblies total, (*17, 67*)), (2) the species’ protein-coding gene sequences were present in the TOGA gene alignment set described in (*14*) (427 species total), and (3) the species was a member of the Boreoeutheria clade (Fig. 1A), given that our primary data were restricted to species within this Magnorder. Although, in many studies, vocal production learning is treated as a binary (“presence / absence”) trait possessed by humans, bats, pinnipeds, and cetaceans alone among boreoeutherian mammals, ‘gold standard’ tests for the trait have been performed for only a handful of species within these clades, and recent reevaluations of the evidence suggest that a more nuanced coding of this complex trait is needed, involving extensive behavioral testing of a greater taxonomic diversity of species (*2, 31, 68*). To attempt to account for this, boreoeutherian species were coded as vocal learners only if they belonged to an established vocal learning clade (i.e, bats, pinnipeds, cetaceans, and humans) and presented evidence of song usage or a rich social acoustic repertoire in the literature. Species potentially falling somewhere outside a simplified binary coding were excluded from analyses based on the following criteria: (i) species with suggestive, insufficient, or controversial evidence of vocal plasticity or learning (*Callithrix jacchus*, *Heterocephalus glaber*, *Indri indri*, *Pan paniscus*, *Pan troglodytes*) or (ii) domesticated species (or subspecies) given demonstrated connection between domestication and increased vocal variability (*69*) (*Camelus bactrianus*, *Camelus dromedarius*, *Canis lupus familiaris*, *Capra hircus*, *Equus asinus asinus*, *Equus caballus*, *Felis catus*, *Mustela putorius furo*, *Vicugna pacos*). In sum, this totaled a set of 175 species with genome assemblies, aligned protein predictions, and vocal learning annotations (25 high-confidence vocal learners and 150 high-confidence non- or very limited-learners, Data S6).

### Evaluating the relationship between relative evolutionary rate of protein-coding genes and vocal learning

We used RERConverge (*16, 46, 70*) to evaluate the relationship between relative evolution rates (RERs) of protein-coding genes and the vocal learning trait across mammals. We obtained a set of 37,552 high-quality protein-coding amino acid alignments generated with TOGA using human reference sequences mapped across a human-referenced MAF alignment of 427 species (*14*). We subsequently filtered these alignments to remove duplicated species, poorly represented proteins, and low-scoring alignments. Specifically, we excluded alignments with fewer than 221 unique species (0.025 quantile of the distribution of unique species number for all alignments), alignments with fewer than 189 total species with ungapped coverage of 50% of the total alignment length (0.1 quantile), alignments with more than 97 duplicated species (0.95 quantile), and alignments with ungapped length <267 bp (0.01 quantile). In total, this resulted in excluding 4,723 transcripts representing 2,613 unique genes. Within the remaining alignments, we excluded any sequences that did not cover 50% of the total alignment length, and, when there were multiple sequences for a species within an alignment, we retained the sequence with the highest identity to the human reference sequence across the full alignment length. For genes with multiple transcripts, we retained only the alignment with the longest median ungapped coverage. From this set of alignments, we then estimated branch lengths on the consensus species tree from (*14*) for each gene using approximate maximum likelihood estimation with the WAG substitution model, as implemented in the phangorn package in R (*71, 72*). We generated RERs for the remaining 16,209 protein-coding genes using RERConverge version 0.3.0 (*16*) with R version 4.1.0. RER-to-phenotype correlations were generated using the vocal learning trait annotations described above and RERconverge’s correlateWithBinaryPhenotype tool, considering all foreground branches and otherwise using default parameters. We corrected the p-values using the Benjamini-Hochberg procedure (*73*) and considered protein-coding genes to have significant associations if their corrected p-values were less than 0.05. In order to exclude the possibility that speciose foreground clades (i.e. bats) were driving our results, we conducted a series of subtree analyses, separately dropping each of the 4 vocal learning clades (bats, cetaceans, pinnipeds, and human) and rerunning the analyses with the remaining 3 clades set as the foreground, identifying the consensus set of convergently accelerated genes robust to all individual clade dropouts (Data S1).

### Subjects for tracing and electrophysiological experiments

All experimental procedures were approved by the Institutional Animal Care and Use Committee of the University of California, Berkeley. All animals were adult Egyptian fruit bats (weight range 130-200g) maintained within the lab colony. For tracing experiments using channelrhodopsin injected into ofM1, subjects were three females born in the lab. For tracing experiments using AAV_DJ_-hSyn-Synaptophysin as an anterograde tracer injected into ofM1, subjects were two males born in the lab. For the 3D mapping of ofM1 descending tracts using dextran amine (Supplementary Figure 1A-B, F and Supplementary Video), the subjects were two females born in the lab. For electrophysiological recordings into ofM1, the four implanted subjects were wild-caught bats. 10 other adult bats (seven males and three females) were used as companions during the recording sessions to elicit vocal interactions. While the age of the four wild-caught bats could not be precisely estimated, all were greater than four years old. Prior to the start of experiments, bats were housed in a communal vivarium. After implantation for electrophysiology, the four implanted bats were initially single housed and subsequently, following recovery from surgery, were co-housed in pairs. All cages were kept in a humidity- and temperature-controlled room, on a 12-hour reversed light-dark cycle. Bats were given *ad libitum* access to water and fed with fruit mix supplemented with honey and vitamins.

### Anesthetic procedure for tracer injection and electrophysiological implant

The general anesthesia and surgical procedures used for Egyptian fruit bats were previously described (*74, 75*). Anesthesia was induced using a cocktail of ketamine, demedotomidine, and midazolam. The depth of anesthesia was evaluated by monitoring the breathing rate, body temperature, and toe pinch reflex. The body temperature was monitored continuously using a rectal probe and maintained around 34.5°C with a heating pad under the bat. If necessary to maintain anesthesia longer than one hour, the bat was injected with a cocktail of dexmedetomidine, midazolam, and fentanyl. Anesthesia was reversed with an injection of atipamezole and the bat was hydrated subcutaneously with lactated ringer’s solution. For all surgical procedures, antibiotics were given for one week post-surgery and analgesic pain medication was given for three days post-surgery. For tracing experiments, the sutures were removed within five days post-surgery.

### Anterograde tracer injection procedures

The procedures for the injection of the anterograde tracer into the brain and retrograde tracer into the cricothyroid muscle were performed at separate times to allow for optimal expression of both tracers (*76*). Two different tracers were used in order to validate the anatomical results across multiple techniques: AAV mediated channel-rhodopsin (ChR2) conjugated to GFP (rAAV5/CamkII-hChR2(H134R)-EYFP, Lot#AV4316LM; UNC Vector Core, NC) (*77*) and a custom synaptophysin/synapsin virus (SYN) from Byungkook Lim’s lab that simultaneously labeled fibers in eGFP and boutons in mRuby2 (AAV_DJ_-hsyn-mRuby2-T2S-Synap-eGFP; Lim Lab, UCSD) (*78, 79*). Each anterograde tracer type was injected bilaterally into ofM1. The 3D reconstruction of ofM1 projections and the localization of the decussation of the pyramidal tract (Supp Fig 1 A-B and Supp Fig 1 F) were obtained in two females injected bilaterally in ofM1 with Dextran amine conjugated to Alexa Fluor 555 (ThermoFisher Scientific; D34679).

For anterograde tracers injected intracranially into ofM1, bats were anesthetized and head fixed in a stereotaxic device (Model 942; Kopf, CA). After opening the scalp with a scalpel, the tissue was retracted to expose the skull. The center of three injection coordinates for ofM1 (AP: +10.72mm, 10.22mm, 9.72mm; ML:+/- 3.2mm) were bilaterally measured from a common reference point above the confluence of the sinus. A small craniotomy (1.2 mm long x 0.6mm wide) was made above ofM1 to expose the surface of the brain while leaving the dura intact. Bilateral injections were made along the anterior-posterior axis into each hemisphere using a NanoFil syringe (36GA beveled needle; WPI, FL) attached to the stereotaxic device. The syringe was slowly lowered to -1.2mm below the surface of the brain around layers V/VI of ofM1 and allowed it to rest for three minutes above the deep target. After pausing, 0.5**μ**L of one of the two anterograde tracers (ChR2, or SYN) were injected at a rate of 4nl/sec using a microinjection pump (UMP3; WPI, FL). The needle was left in place for five minutes at each site. A total volume of 1.5**μ**L was delivered in each hemisphere. Upon completion of the six injections, Kwik-Sil (WPI, FL) was used to fill the craniotomy and protect the brain and the tissue was sutured.

### Retrograde injections into cricothyroid muscles

The retrograde tracer injection was performed approximately one month following the anterograde tracer injection to optimize maximal expression of the virus and propagation of cholera toxin B (CTB). Approximately one week before the planned perfusion time, the bats were anesthetized according to the same procedures above. Once anesthetized, the neck was shaved and the bat was placed on its back on a heating pad to facilitate access to the larynx. The skin overlying the larynx was incised using a scalpel to reveal the larynx below the sternohyoid and infrahyoid muscles. The tissue was retracted to expose the cricothyroid muscle caudal to the inferior border of the thyroid cartilage and medial to the cricothyroid joint.

Bilateral injections of cholera toxin B conjugated to fluorescent labels (ThermoFisher, C34778, AlexaFluor 647) were made in the cricothyroid muscle at six different points, three on each side. A NanoFil syringe (35GA beveled needle; WPI, FL) attached to the stereotaxic device was slowly lowered approximately -0.4mm below the surface of the muscle. After waiting one minute to allow the tissue to settle, 2**μ**L of CTB was injected at a rate of 16nl/sec into each injection site. The needle was left in place before moving to the next injection site. Upon completion of the six injections of 2**μ**L the tissue was sutured and two surgical staples were placed over the sutures.

### Electrophysiological implant surgery

The surgical procedures for the implantation of four tetrode microdrives followed those described previously in detail for Egyptian fruit bats (*26, 75, 80*). Each bat was implanted with a lightweight microdrive (Harlan 4-Drive, Neuralynx; weight 2.1 g) loaded with four tetrodes (∼45 **μ**m diameter; four strands of platinum-iridium wire, 17.8 **μ**m in diameter, HML-insulated) that could be moved independently. The tetrodes exited the microdrive through a guide cannula with ∼300 **μ**m horizontal spacing between tetrodes. On the day before surgery, each tetrode’s tip was cut flat using high-quality scissors (tungsten-carbide scissors ceramic coating, CeramaCut; FST) and plated with Gold Plating Solution (Neuralynx) to reduce the impedance of individual wires to 0.20-0.55 MΩ (at 1 kHz). The principal surgical steps to implant the microdrive were the following: after scoring the skull to improve adhesion and mechanical stability, a circular craniotomy of 1.7 mm diameter was made in the skull over the left hemisphere 3.2 mm lateral to the midline and 10.7 mm anterior to the transverse sinus that runs between the posterior part of the cortex and the cerebellum; after a durotomy, the microdrive was placed vertically such that the tip of the microdrive’s guide tube was flush with the brain’s surface; the exposed craniotomy was then filled with a biocompatible elastomer (Kwik-Sil, World Precision Instruments) to protect the brain; a bone screw (FST) with a soldered stainless-steel wire was fixed to the skull in the frontal plate contralateral to the microdrive to serve as a ground screw; an additional set of 3 bone screws were fixed to the skull to serve as anchor screws for maintaining mechanical stability of the implant; finally the exposed skull and anchor screws were covered with a thin layer of quick adhesive cement (C&B Metabond, Parkell) to provide a substrate for the adhesion of dental acrylic that was added as final layer to secure the entire microdrive to the screws and to the skull.

### Electrophysiological and audio recording devices

Electrophysiological recordings were conducted using a wireless neural data logging device (“neurologgers”; MouseLog-16 (vertical version), Deuteron Technologies) that interfaced with the microdrive of each implanted animal. The neurologger amplified the voltage signals from the 16 channels of the four tetrodes referenced to the ground screw, performed analog-to-digital conversion at a sampling rate of 31.25 kHz, and stored the digitized data on a removable 32GB SD card that can hold up to 9 hours of recording. The system has a bandwidth of 1 Hz - 7 kHz, records voltage with a fine resolution of 3.3 μv, and has a low level of noise generally close to the limit of Johnson noise from the impedance of a given source. The neurologger and its lithium-polymer battery were encapsulated in a house-made 3D-printed plastic casing to prevent damage to the electronics, and weighed a total of 9.9 g. The audio recordings of each individual bat vocalizations were performed using a call detector, as previously described (*26*). In brief, a single-axis, low mass, piezo-ceramic accelerometers (BU-27135, Knowles Electronics, sensitivity 0-10kHz) was mounted on a flexible rubber necklace placed against the throat of the subject in a way that did not restrict normal behavior to detect laryngeal vibrations produced during vocalizations. The signal of the accelerometer was recorded, digitized at a sampling rate of 50Hz, and saved on removable SD cards with a wireless audio data logging device (“audiologgers”; Audio Logger AL1, Deuteron Technologies) mounted on the necklace on the back of the subject. The audiologger and its lithium-polymer battery were encapsulated in a house-made 3D-printed plastic casing to prevent damage to the electronics. All audiologgers and neurologgers were controlled and synchronized by a single transceiver. The Egyptian fruit bats in our experiment weighed more than 140gr and could fly with ease while equipped with both the neurologger and the audiologger.

### Vocalizations and motor actions recording sessions

The four implanted bats were divided into two pairs that were independently recorded for 1-3 hours per day over multiple days (16 and 32 sessions) with 2 or 3 peers. These 2-3 peers were randomly chosen from a pool of ten bats (7 males, 3 females) and were used as companions to increase the probability of vocal interactions implicating the subjects. During the daily electrophysiological and audio recording sessions these groups of 4 to 5 bats (2 implanted + 2-3 companions) were housed in a rectangular prism (180 x 60 x 60 cm) that had two sides made of plexiglass, thereby permitting clear remote visual monitoring of bats behavior via 2 cameras (Flea 3 FLIR). The remaining sides of the enclosure were made of plastic mesh, allowing bats to easily perch and crawl on the surface. The enclosure was placed in an electromagnetically and acoustically shielded room and all recording sessions were conducted during the dark cycle under red LED light. All bats were equipped with audiologgers such as to record and identify their vocalizations. Water was given ad libitum and fruits were placed into the cage such as to engage the animals into chewing and licking behavior. The experimenter was monitoring the behavior of the animals in an ante-chamber via video cameras and an ambient ultrasonic microphone (Earthworks, M50) centered 20cm above the cage ceiling and connected to the main computer unit via an analog to digital convertor (MOTU, 896mk3). Using a house-made keystroke annotation code written under Matlab, the experimenter was manually annotating chewing, licking (self-grooming using licking movements), and quiet (staying still in a relaxed position, wing closed) behaviors. The audio was recorded throughout the session (sampling rate of 192kHz) from the ambient microphone using an in-house Matlab GUI (VocOperant; https://github.com/julieelie/operant_bats). The synchronization between the microphone recording, the manual annotation of behaviors and the transceiver controlling the audiologgers and neurologgers was achieved using transistor-transistor logic (TTL) pulses generated by an UltraSoundGate Player 216H (Avisoft Bioacoustics) and sent via coaxial cables. After each recording session, tetrodes were connected to a wired recording system (Digital Lynx, Neuralynx) to monitor the neural signals and advance the tetrodes. Tetrodes were moved downward once every one to two days (generally by 100**μ**m) in order to record single units at new sites.

### Histology

#### Perfusion

Approximately one week following the injection of the retrograde CTB tracer, or after the last day of electrophysiological recording, bats were administered an overdose of pentobarbital and perfused transcardially with 250ml of pH 7.4 phosphate buffered saline (PBS) spiked with 0.5ml heparin (1000 USP units/ml) •••followed by 250ml of fixative (3.7% formaldehyde in phosphate buffered saline). When the brain was implanted with electrodes, tetrodes were left in the brain for 30 minutes before extracting them. The brain was then carefully removed from the skull and post-fixed overnight in the same fixative. To avoid over-fixation, the brain was removed from fixative after 24 hours and switched into a 30% sucrose solution for cryoprotection. After approximately two days or once the brain had sunk to the bottom, 40**μ**m coronal sections were made using a microtome (HM450; ThermoFisher, MA) with a freezing stage.

#### Staining and Immunocytochemistry

The sections from the implanted bats were Nissl-stained with cresyl violet. Slides were imaged using a light microscope to verify tetrode positions.

The sections of the brains from bats injected with tracers were stained for VGLUT1 using a fresh tissue floating immunohistochemistry protocol. Immunohistochemistry was conducted in 12-well plates filled with 4 ml of solution for washes and blockings and 48 well plates filled with 1 ml of solution for primary and secondary incubation. Fifteen brainstem slices were selected from each series centered around the NA region and three brainstem slices anterior to the target region were selected for antibody control staining. Briefly, the tissue was placed in floating wells on a lab rotator in a cold room at 4°C and washed in three separate PBS (0.025M, pH 7.4) solutions for five minutes each wash. The tissue was then moved to a blocking solution containing 10% goat serum (Sigma-Aldrich, G9023) in 0.3% triton-PBS (Triton X-100 - ACROS Organics, 21568-2500 in 0.025M PBS) and rotated for 90 minutes at 4°C. The tissue was incubated overnight at 4°C in rabbit anti-VGLUT1 primary antibody (provided by Eiman Azim, Salk Institute for Biological Studies and produced in Tom Jessell’s lab at Columbia University) (*81*), which was prepared in a 1:16,000 dilution in 5% goat serum and 0.3% triton-PBS. Control slices were incubated in an antibody buffer without primary antibody. Approximately 16-24 hours after the start of the primary antibody incubation, the tissue was moved into three separate 0.3% triton-PBS washes for 10 minutes in each wash at 4℃. The secondary antibody was goat anti-rabbit conjugated to a fluorescent protein (ThermoFisher A27012, A27018) that did not conflict with the anterograde or retrograde tracers (selected wavelength 594 nm for bats injected with ChR2). The tissue was incubated in the secondary solution diluted 1:500 in 5% goat serum and 0.3% triton-PBS at room temperature for 90 minutes before three final washes in PBS (0.025M, pH 7.4) for 10 minutes each. DAPI was added to the secondary solution for the final 10 minutes of incubation at 1:10,000 dilution (D1306; ThermoFisher, MA). The sections were then mounted on glass slides and cover-slipped using ProLong Gold Antifade Mountant (P36934; ThermoFisher, MA).

### Imaging and anatomical quantification

#### Fluorescent imaging

All imaging was conducted at the University of California, Berkeley Cancer Research Laboratory Molecular Imaging Center and the Henry H. Wheeler Jr. Brain Imaging Center at UC Berkeley. Preliminary imaging at magnification of 10x/20x using Plan-Apochromat 10x/20x objective was conducted on Zeiss Axio Scan Z1 Slide Scanner. Slices were imaged in four fluorescent channels – AF647 (Excitation: 653 nm, Emission: 668 nm, Gain: 0, Exposure time: 25ms, Filter Cube: 50 Cy5), AF488 (Excitation: 493 nm, Emission: 517 nm, Gain: 0, Exposure time: 20ms, Filter Cube: 38 HE Green Fluorescent Protein), DAPI (Excitation: 353 nm, Emission: 465nm, Gain: 0, Exposure time: 5ms, Filter Cube: 49 DAPI), and AF598 (Excitation: 570 nm, Emission: 618 nm, Gain: 0, Exposure time: 15 ms, Filter Cube: 64 HE mPlum). All images were taken with 100% fluorescent lamp strength on Hamamatsu Orca Flash Camera with 1x Camera Adapter.

More specific imaging of nucleus ambiguus area and other target and control regions was conducted at magnification of 63x with Plan Apochromat 63x/1.4 Oil DIC M27 Objective with target area (135 **μ**m x 135 **μ**m) on Zeiss LSM 880 Confocal Microscope at the UC Berkeley Molecular Imaging Center. The target region was localized by finding retrogradely labeled cells, centering them, and taking a z-stack to encompass the entire cell volume. Each z-stack was taken over 1.5 **μ**m depth with the above mentioned area and 1 **μ**m between each z-stack slice with 4 fluorescent channels – AF 647 (Laser wavelength: 633 nm, Excitation: 633 nm, Emission: 697 nm, Laser Power: 6.0%, Detector Gain: 650, Detector Digital Gain: 1, Detector Offset: 0), AF 594 (Laser wavelength: 594 nm, Excitation: 594 nm, Emission: 659 nm, Laser Power: 8.0%, Detector Gain: 675, Detector Digital Gain: 1, Detector Offset: 0), AF 488 (Laser wavelength: 488 nm, Excitation: 488 nm, Emission: 552 nm, Laser Power: 5.0%, Detector Gain: 625, Detector Digital Gain: 1, Detector Offset: 0), and DAPI (Laser wavelength: 405nm, Excitation: 405nm, Emission: 462nm, Laser Power: 6.0%, Detector Gain: 625, Detector Digital Gain: 1, Detector Offset: 0). If relevant, AF 561 (Laser wavelength: 561 nm, Excitation: 561 nm, Emission: 621 nm, Laser Power: 8.0%, Detector Gain: 650, Detector Digital Gain: 1, Detector Offset: 0) was taken on a separate z-stack due to maximum channels/image microscope limitations. All confocal images were taken as 12-bit images with line-by-line imaging and 2x averaging.

Deconvolution was conducted using HuygensPro software from the Biological Imaging Facility at UC Berkeley. Theoretical PSF was generated using LSM 880 confocal microscope and 63x objective parameters. Image histograms were created using a logarithmic mapping function and background was generated using automatic estimation with an area radius of 0.7 microns. Deconvolution was conducted for each channel with 200 maximum iterations, 30 signal to noise ratio, 0.01 quality threshold, and with optimized iteration mode.

#### 3D model of ofM1 tracing and localization of decussation

Series of coronal slices were manually stacked to build a 3D model of the fluorescent pathways of fluorescently-labeled dextran amine (DA) tracing to/from ofM1. Bats were injected with fluorescent dextran amine in ofM1 and, following perfusion, coronal slices of the whole brain separated by 200 **μ**m were stained for DAPI to create a uniform background marker. After slide scanning the entire brain (see scanning settings above), the color levels for the DAPI and DA channels were equalized for every coronal slice and exported into lossless tiffs from ZEISS ZEN Microscope Software. The coronal slices were cut out from against the background and manually aligned using Adobe Photoshop so that the edges lined up in a full stack from the rostral tip of the olfactory bulbs to the caudal tip of the spinal cord. All images were then stacked in Imaris to create a 3D model of the brain that can be rotated to observe the whole structure of the corticobulbar pathway from ofM1 to NA (Supplementary Figure 1A-B and Supplementary Video).

#### Image quantification

Total numbers of cells, fibers, boutons, and DAPI-labeled cells of brains injected with SYN and ChR2 were quantified using Imaris software (Version 9.2.1) at the UC Berkeley Cancer Research Laboratory Molecular Imaging Center. Retrogradely labeled cells and boutons were automatically counted using Imaris Spot Tracker. Initially, the diameter of retrogradely labeled cells (15-30 **μ**m), boutons (1-3 **μ**m), and DAPI labeled nuclei (10-15 **μ**m) was measured using automatic measuring tools in Imaris in a 2D slice and imputed as a parameter for Spot Tracker. After computing, x, y, and z, diameter and position was manually adjusted until the spot covered the entire retrogradely labeled cell. Fibers were counted and traced using semi-automatic Imaris AutoPath Tracer by manually tracing along the length of the fiber. Fiber diameter was set to 1.4 **μ**m across all z-stacks.

Overlap was quantified manually by determining points of colocalization between fiber/cell and fiber/cell/bouton. Z-stacks were exported as single multi-channel TIFF images and then opened each Z-stack individually in Adobe Photoshop. Points of co-localization were individually and manually marked. TIFF images were compared to the original z-stack to confirm the existence of fiber/cell/bouton/DAPI-labeled cells over multiple slices and ensure real signal. The number of retrogradely labeled cells with CTB that had at least one overlap with fibers and the number of retrogradely labeled cells with CTB that had at least one overlap with fibers and boutons were counted. If overlap on the same cell spanned multiple TIFF images, it was only counted in the first slice in which it appeared.

### Acoustic data processing

Acoustic data logged as voltage traces on the SD cards of the audiologgers were extracted into Matlab files and aligned across bats simultaneously recorded using a custom-made Matlab code (https://github.com/NeuroBatLab/LoggerDataProcessing/). Potential vocalizations were then detected and segmented from these piezo recordings using an in-house series of Matlab scripts (https://github.com/NeuroBatLab/SoundAnalysisBats). The whole process consisted in three major steps: 1) detection of sound events on call detector signals, 2) automatic classification between vocalizations and noise, 3) manual curation of potential vocalizations.

1. Detection: To focus on the detection of vocalizations emitted by the bat wearing the collar, the signal of each call-detector was first band passed between 1 and 5kHz. As previously described, this frequency range is not contaminated by airborne vocalizations from other bats standing close to the collar wearer (*26*). After determining a noise threshold from sections of silence during the recording session for that call-detector, potential vocalizations were detected by threshold crossing on the amplitude envelope (RMS, sampling frequency 1kHz) and any sound event above threshold for longer than 7ms was kept.
2. Automatic step of data sorting between actual vocalizations and noise: Sound events closer than 50ms were merged as a single sound sequence and a battery of 20 acoustic measurements were applied on them (see (*82*) for mathematical definitions): RMS, maximum amplitude, the five first momentum of the amplitude envelope taken as a distribution (mean, standard deviation, kurtosis, skewness, and entropy), the five first momentum of the frequency spectrum taken as a distribution, the three quartiles of the frequency spectrum, the mean pitch saliency, and four parameters that pertain to the sound as recorded from the ambient microphone (RMS, mean and maximum of the amplitude envelope, and maximum value of cross-correlation between the microphone and the call detector signals). Potential vocalizations among the detected sound events were then identified using these 20 acoustic parameters as input to a support vector machine trained on the data of two sessions manually sorted between vocalizations and noise by an expert (JEE). This automatic sorting was set to be very conservative of vocalizations by setting the threshold on the posterior probability of a vocalization at 0.2.
3. Manual curation: to further eliminate noise and check the identity of the bat producing vocalizations, each potential vocalization was aurally and visually scrutinized by an expert (JEE) based on the inspection of the spectrograms of its signal as recorded from the ambient microphone and from the call detectors of all the bats. After this step, vocalizations further than 200ms apart were considered as distinct, while the others were merged into a single sequence.

### Neural data processing

#### Spike detection and sorting

Neural data logged as voltage traces on the SD cards of the neurologgers were extracted into Matlab files and aligned across simultaneously recorded bats and with the audio data, using custom Matlab code (https://github.com/NeuroBatLab/LoggerDataProcessing/). Spike detection and sorting was achieved using a generative algorithm (Kilosort 2.0, https://github.com/MouseLand/Kilosort/releases/tag/v2.0, (*83*)) with the parameters set as indicated under https://github.com/julieelie/Kilosort2_Tetrode/configFiles/configFile16.m. Clusters were further manually curated using Phy (https://github.com/cortex-lab/phy).

Units obtained after manual curation were sorted between multi-units and single units by applying thresholds on two metrics: 1) the consistency of the spike snippet (signal over noise ratio of the spike snippet, SNR, estimated at the four contact points of the tetrode) and 2) the respect of the refractory period (percent of violation of the refractory period, VRP). A unit had to have at least one out of four SNR values above 5 and VRP<0.1% to be considered as a single unit.

For each unit, the spike snippet SNR was calculated for each electrode of the tetrode as:

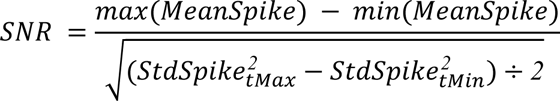

with *MeanSpike* the average spike snippet over all spikes, *stdSpike* the time-varying standard deviation of the spike snippet over all spikes, and *tMax* and *tMin* the time point at which the average spike snippet reaches maximum and minimum values, respectively.

Applying the above mentioned thresholds on SNR and VRP of units manually curated with Phy yielded 381 single units (SNR = 8.15 +/- 0.12; VRP = 0.0308 +/- 0.0015 %). 94.2% of single units (359/381) had an index of contamination Q (*83–85*) below 0.2 (Q = 0.0531 +/- 0.0042, N=381) and a significant test of refractoriness against a Poisson distribution at p<0.05.

#### Firing rate calculations and analysis

For this analysis, we selected the single units that were recorded during the sessions when the subject had produced a minimum of ten vocalizations longer than 100 ms and had displayed chewing, licking and quiet behaviors (n= 237 units). For each single unit, the time average firing rate during the production of vocalizations longer than 100ms was estimated for the duration *D* of each vocalization starting 10ms before vocalization onset as:

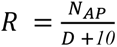 with *N*_*AP*_ the number of action potentials occurring during the time window *D* + *10 ms*.

For all other behaviors (chewing, licking, and quiet), the annotated period of time where the bat was demonstrating that behavior was reduced by 1 second (true offset considered as 1s before keystroke) to conservatively accomodate for the annotator time response. Then the firing rate was estimated in time segments of the same durations as those used to estimate firing rate during vocalizations by randomly sampling without replacement into the period of time where the bat was demonstrating the behavior of interest.

Firing rate comparisons between pairs of behaviors were achieved by applying a test on the deviance of the Poisson Generalized Linear Model (GLM) predicting the rate *R* as a function of the category of behavior (function fitglm of Matlab). P-values were corrected for false detection rate using the Benjamini-Hochberg procedure.

#### Information on coherence during vocalization perception and production

The relationships between each single unit activity and vocal production or vocalization perception were quantified by calculating the coherence between the time varying amplitude of vocalizations and the time varying firing rate of the unit during the vocalizations respectively produced or heard. For this analysis, we selected the single units that were recorded during the sessions when the subject had both produced and heard a minimum of twenty vocalizations (n=219 units). Vocal activity, as measured by the piezoelectric sensor of the call detector, and neural activity, as represented by the arrival times of action potentials, were collected from 200ms prior to the onset to 200ms after the offset of each vocalization. The time-varying amplitude of each sound extract was taken as the amplitude envelope calculated with a frequency cut-off at 150 Hz and sampled at 1000 Hz (BioSound python package, https://github.com/theunissenlab/soundsig, (*82*)). The spike pattern corresponding to each sound extract was convolved with a Gaussian window of 1ms standard deviation to obtain a time-varying firing rate sampled at 500 Hz. The coherence between the time-varying amplitude and the time-varying firing rate was then calculated across all vocalizations that the animal had produced/heard during the session to give the motor/auditory coherence. A multitaper approach was implemented in the estimation of coherence to obtain an error measure (*86*). The information on coherence was obtained by integrating all values of positive coherence up to the Nyquist limit. Information was calculated on the estimate of coherence and on its lower and upper bounds both for vocalizations produced and vocalizations heard, yielding a value of motor information *Info*_*motor*_ and its corresponding lower and upper bounds 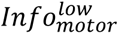 and 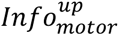 as well as a value of auditory information *Info*_*aud*_ and its lower and upper bounds 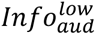 and 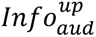 For each unit, the information D-Prime was then calculated as:

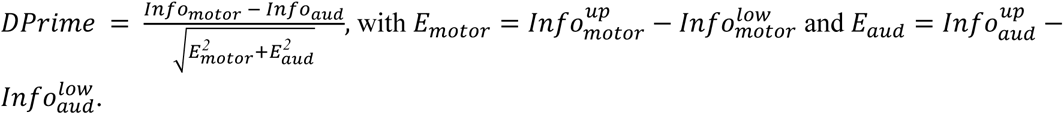

The significance of the difference in information between produced (motor) and heard (auditory) vocalization was assessed by a likelihood ratio test between two linear mixed effect (LME) models predicting the information value with or without the type of information (motor vs auditory) as a fixed effect and with the identity of the subject as a random variable.

### Animals and sample collection for epigenomics

All animal procedures were in accordance with the National Institutes of Health Guide for the Care and Use of Laboratory Animals and approved by the Institutional Animal Care and Use Committees the University of California, Berkeley. Two adult (>1 year) male Egyptian fruit bats (*Rousettus aegyptiacus*), one male and one female, were housed socially in a large free-flight vivarium. Bats were acoustically and socially isolated in a sound recording chamber (*external dimensions*: 61 cm X 65 cm X 61 cm; *internal dimensions*: 51 cm X 61 cm X 61 cm) the night prior to experiments. Bats were acoustically monitored to confirm non-vocalizing status prior to experiments in order to control for the effects of activity-induced expression. To control for circadian effects, all experiments were performed between 8 and 10am. Bats were administered with an overdose of pentobarbital with an intraperitoneal injection. We then rapidly opened the skull and removed the brain using round-tipped safety scissors. Brains were sliced coronally into 300 μm sections in a vibrating microtome (Leica VT 1200) in ice-cold, oxygenated artificial cerebrospinal fluid [119 mM NaCl, 2.5 mM KCl, 1 mM NaH2PO4 (monobasic), 26.2 mM NaHCO3, 11 mM glucose] and regions of interest were excised under a dissection microscope. Liver, muscle, and gonads were collected immediately. Tissues were preserved in a cryoprotectant medium (CryoStor CS10, Biolife Solutions) in cryovials, which we placed in a foam freezing container (CoolCell, Corning) and transferred to a -80°C freezer in order to ensure a controlled freezing rate of -1°C per minute.

### CryoATAC-seq protocol

Tissue samples were processed as described previously (*42, 87, 88*) with the following minor differences in procedure and reagents. Cryopreserved samples were warmed in a 37°C water bath for 2 minutes and then transferred into 12 mL PBS supplemented with a protease inhibitor cocktail (Roche) and gently mixed by inversion. Samples were centrifuged at 300 rcf for 5 minutes at 4°C before aspirating all supernatant and resuspending the samples in ice cold lysis buffer (*42*). Nuclei were isolated from dissected tissues using 30 strokes of homogenization with the loose pestle (0.005 in. clearance) in 5mL of cold lysis buffer placed in a 15 mL glass Dounce homogenizer (Pyrex #7722-15). Nuclei suspensions were filtered through a 70 μm cell strainer, pelleted by centrifugation at 2,000 x g for 10 minutes, resuspended in water, and filtered a final time through a 40 μm cell strainer. Sample aliquots were stained with DAPI (Invitrogen #D1206), and nuclei concentrations were quantified using a manual hemocytometer under a fluorescent microscope. Approximately 50,000 nuclei were input into a 50 μL ATAC-seq tagmentation reaction as described previously (*42, 87*). The resulting libraries were amplified to 1/3 qPCR saturation, and fragment length distributions estimated by the Agilent TapeStation System showed high quality ATAC-seq fragment length periodicity. We shallowly sequenced barcoded ATAC-seq libraries at 1-5 million reads per sample on an Illumina MiSeq and processed individual samples through the ENCODE ATAC-seq pipeline (version 1.8.0, accessed at https://github.com/ENCODE-DCC/atac-seq-pipeline) for initial quality control. We used the QC measures from the pipeline (clear periodicity, library complexity, and minimal bottlenecking) to filter out low-quality samples and re-pooled a balanced library for paired-end deep sequencing on an Illumina NovaSeq 6000 System through Novogene services to target >30 million uniquely mapped fragments per sample after mitochondrial DNA and PCR duplicate removal.

### ATAC-seq data processing

We processed raw FASTQ files of ATAC-seq experiments with the ENCODE ATAC-seq pipeline (version 1.8.0, accessed at https://github.com/ENCODE-DCC/atac-seq-pipeline) to identify open chromatin region (OCR) peaks from sequenced samples. We ran the ENCODE pipeline using the mRouAeg1.p assembly (*89*). We ran the pipeline with the default parameters except for “atac.multimapping” : 0, “atac.cap_num_peak”: 300000, “atac.smooth_win”: 150, “atac.enable_idr”: true, and “atac.idr_thresh”: 0.1. We generated filtered bam files, peak files, and signal tracks for each biological replicate and the pool of replicates for each tissue. To account for differences in sequencing depth between samples, we identified reproducible peak sets, which we defined as peaks with an irreproducible discovery rate (IDR, (*90*)) < 0.1 across pooled pseudo-replicates. We found that all of our samples except for the two liver samples were high-quality, displaying high fragment length periodicity, low PCR duplicate rates, and concordance between biological replicates.

In addition to identifying peak sets for individual tissues, we identified consensus peak sets to serve as genome-wide background sets representing the intersection of the reproducible open chromatin peaks identified from all processed tissues, from all cortical samples, or from all motor cortex samples. We obtained these background sets using bedtools (version 2.25.0, (*91*)) intersect with the -wa and -u options to combine reproducible peak sets. We prepared OCRs for downstream analysis in the following way: we combined peaks within 50 bp of one another using bedtools merge, preserving the summit location as the average of the summits of all merged peaks. We used bedtools subtract with option -A to remove those peaks that were within 1.5 kb from any annotated coding or noncoding exons, enabling us to exclude promoters, coding sequences, and noncoding RNAs from our background set. We further filtered out peaks greater than 1.5 kb in width. In order to identify the complete set of exonic exclusion regions for Egyptian fruit bat, we used the complete set of mRouAeg1.p annotations (*89*).

To identify OCR peaks differentially active between tissues, we first quantified the number of reads from each tissue that aligned to the consensus peaksets described above using featureCounts (*92*). We then contrasted the read counts at each peak between tissues using the negative binomial model in the DESeq2 R package (*93*). We identified differential peaks based on their Wald statistic value, with a statistical cutoff of p < 0.05 (Data S3).

In order to identify OCR orthologs across species, we aligned the consensus peaksets as well as their peak summits across all of the species present in the Zoonomia Cactus alignment (*17, 67*) using halLiftover (*94*) with default parameters. We filtered the raw outputs of halLiftover and assembled them into contiguous OCRs using HALPER (*95*) with parameters -max_frac 2.0, -min_len 50, -protect_dist 5, and - narrowPeak. We obtained the sequences underlying these OCR summit orthologs’ +/- 250bp using fastaFromBed in bedtools (*91*).

### Gene functional enrichment analyses

We performed gene ontology analyses using GREAT version 4.0.4 (*96*) and g:Profiler (*97*). We ran GREAT on the genomic coordinates of ofM1 vs wM1 differential bat OCR peak sets mapped to human (hg38) using halLiftover (*94*) and HALPER (*95*) with the same parameters that were used for other analyses (Data S4). We also ran g:Profiler to identify functional enrichments in genes demonstrating relative evolutionary rate convergence in vocal learners using default parameters (Data S2).

### Transcription factor binding motif enrichment

To identify transcription factor binding motifs enriched in differential OCR peak sets of interest relative to shuffled sequences, we used AME in the MEME suite (*98, 99*), setting all parameters to default. For our position weight matrix set, we used the JASPAR2018 CORE set of non-redundant vertebrate motifs (*100*).

### Predicting OCR ortholog open chromatin activity across species and associating predictions with vocal learning

We used the Tissue-Aware Conservation Inference Toolkit (TACIT) to identify open chromatin regions (OCRs) whose predicted open chromatin differences between species are associated with differences in vocal learning. Specifically, we investigated the association between vocal learning and open chromatin predictions across 222 boreoeutherian species from Zoonomia from models trained on Egyptian fruit bat (this study), Brown Norway rat (*19*), C57BL/6J mouse (*20*), and Rhesus macaque motor cortex (intersection of ofM1 and wM1 peaks, limited to peaks < 1kb, > 20kb from the nearest protein-coding TSS, and not overlapping protein-coding exons) (*19*) as well as human and mouse motor cortical parvalbumin (PV+) neuron (*21, 53*) open chromatin using TACIT, as described in a companion paper to this study (*22*). From the cross-species mapped OCR consensus peak sets, we removed motor cortex OCRs that did not have usable orthologs in at least half of the species with vocal learning annotations (provided in Data S6) and at least three vocal learning species, as these were unlikely to have sufficient power to identify associations. Likewise, we removed motor cortex OCRs that did not have a usable ortholog in at least one species in Chiroptera (bats), the order with the largest number of mammalian vocal learners. In addition, we removed motor cortex OCRs that did not have at least one usable ortholog in a non-chiropteran vocal learner, as any association we found for such OCRs would likely be explainable by phylogeny (i.e., driven by bats) and therefore not be able to be found through phyloglm (*54, 55*), the method that we used for associating predicted open chromatin with phenotypes while correcting for phylogeny. After applying these filtering steps, we corrected our empirical p-values from phylogenetic permulations (*22, 56*) using Benjamini-Hochberg (*73*).

## Supplemental Tables

**Table S1:**
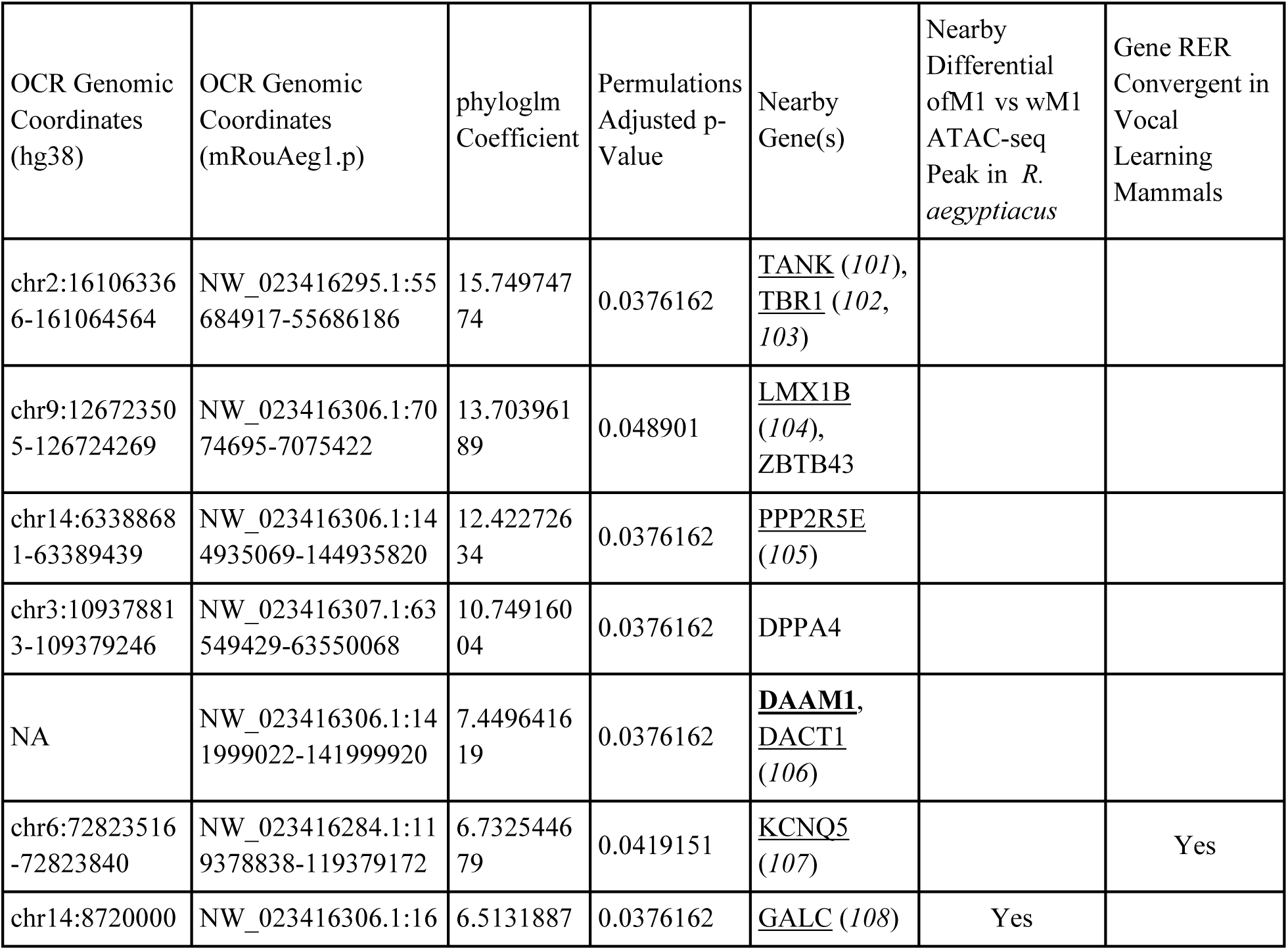
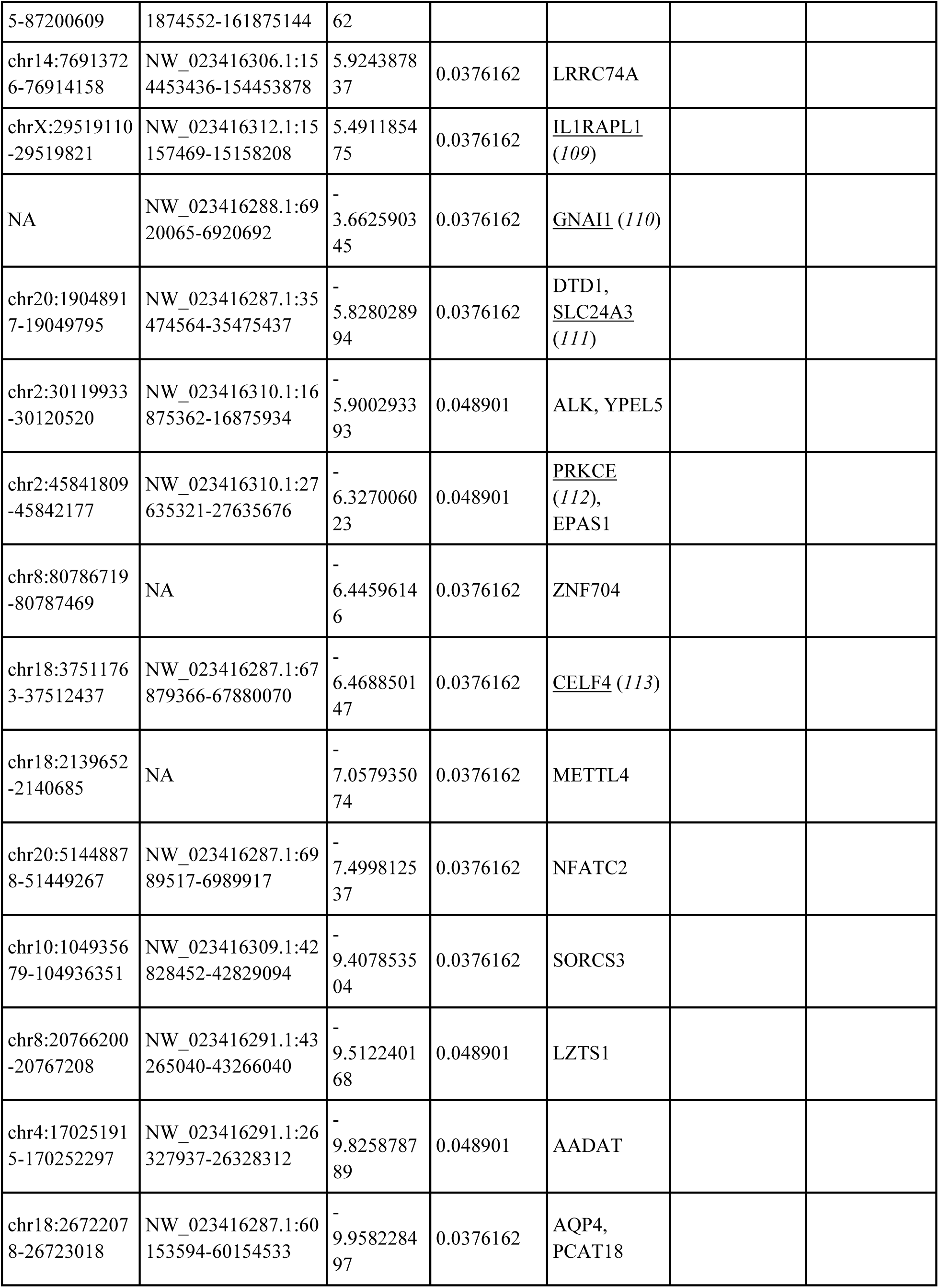
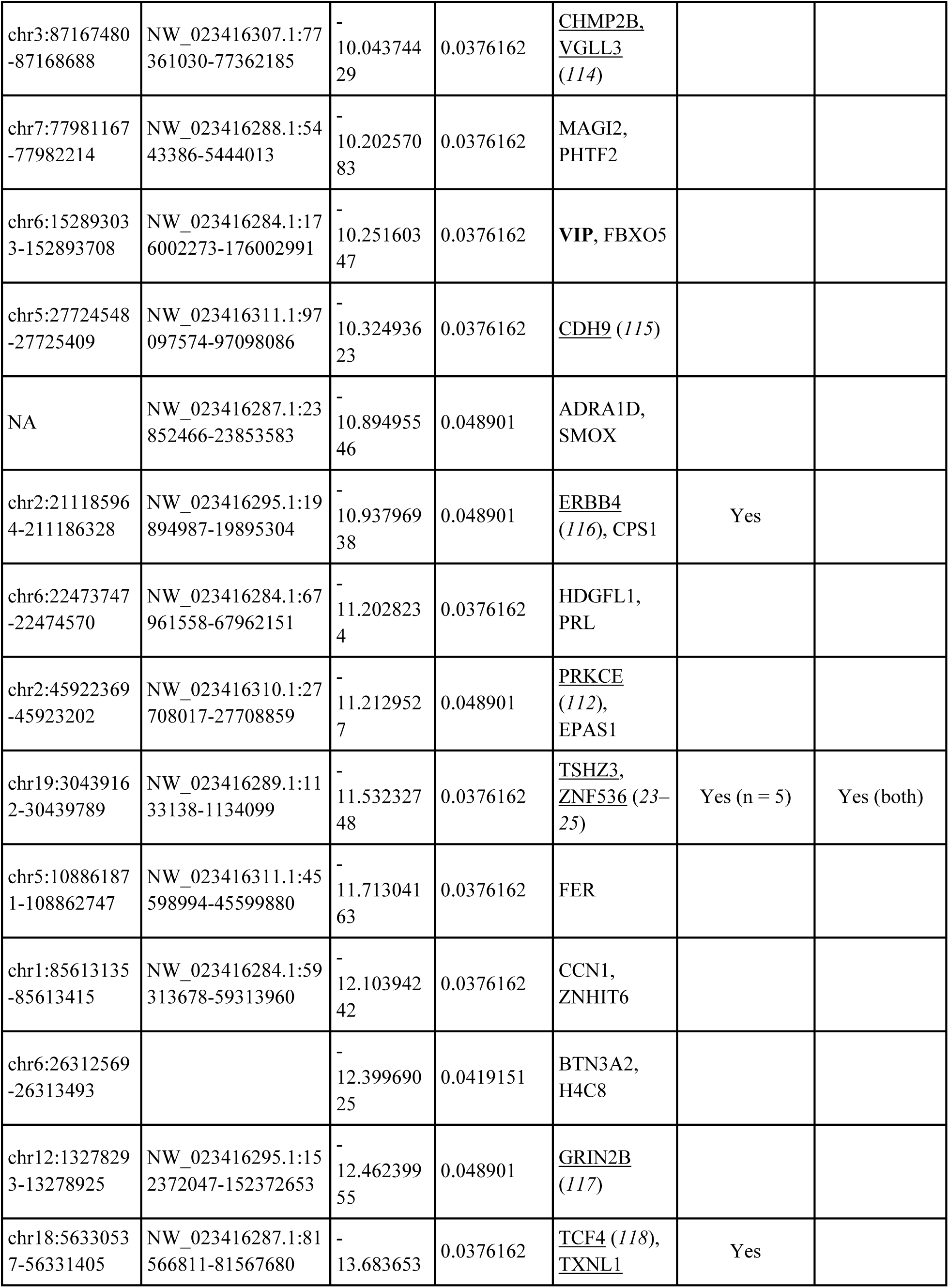
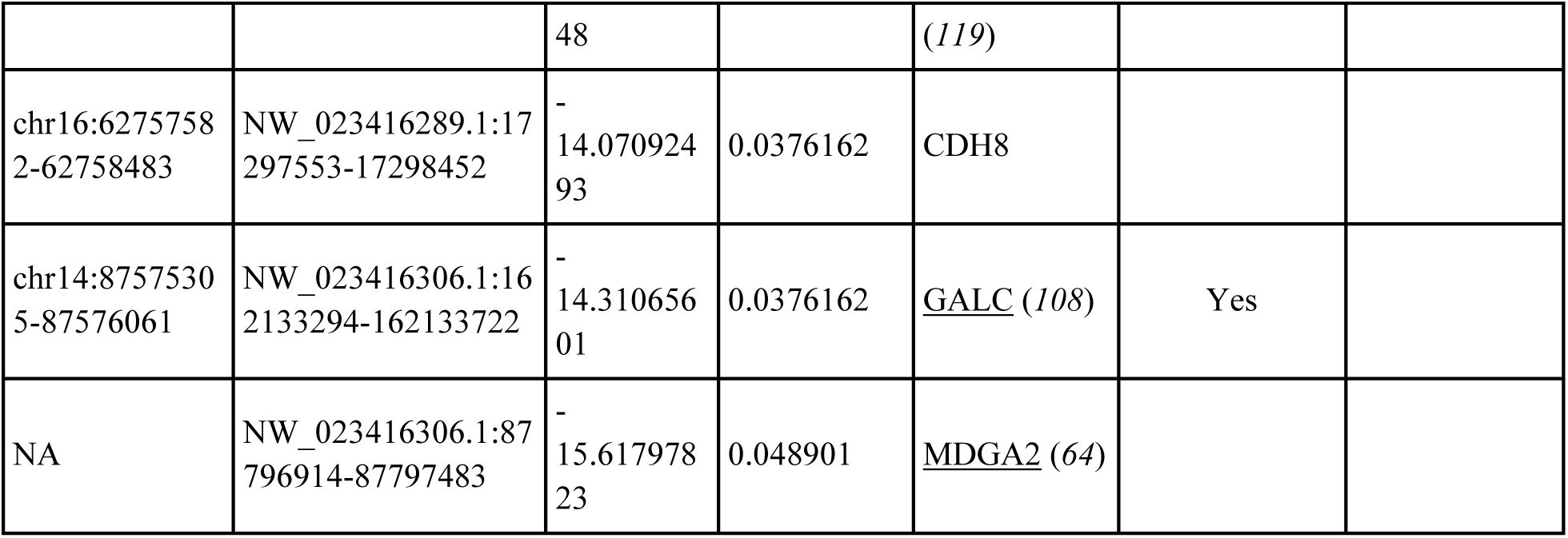
M1 OCRs whose predicted open chromatin state in boreoeutherian mammals is significantly associated with vocal learning. Coordinates are reported for both human (hg38) and *R. aegyptiacus* bat (mRouAeg1.p). The coefficients shown are those reported by phyloglm for the association between the vocal learning trait and the OCR orthologs’ open chromatin prediction. The permulations p-values represent the adjusted p-values after Benjamini-Hochberg correction. Nearest genes are indicated, with genes previously associated (in some cases specifically, in other cases as part of larger deletions) with speech delay or disability underlined (with associated references) and genes previously shown to be convergently regulated in humans and song-learning birds (*4*) in bold. OCRs are ordered by phyloglm coefficient, from high to low (indicating OCRs with higher or lower predicted open chromatin values, respectively, in vocal learners relative to vocal non-learners).

**Table S2:**
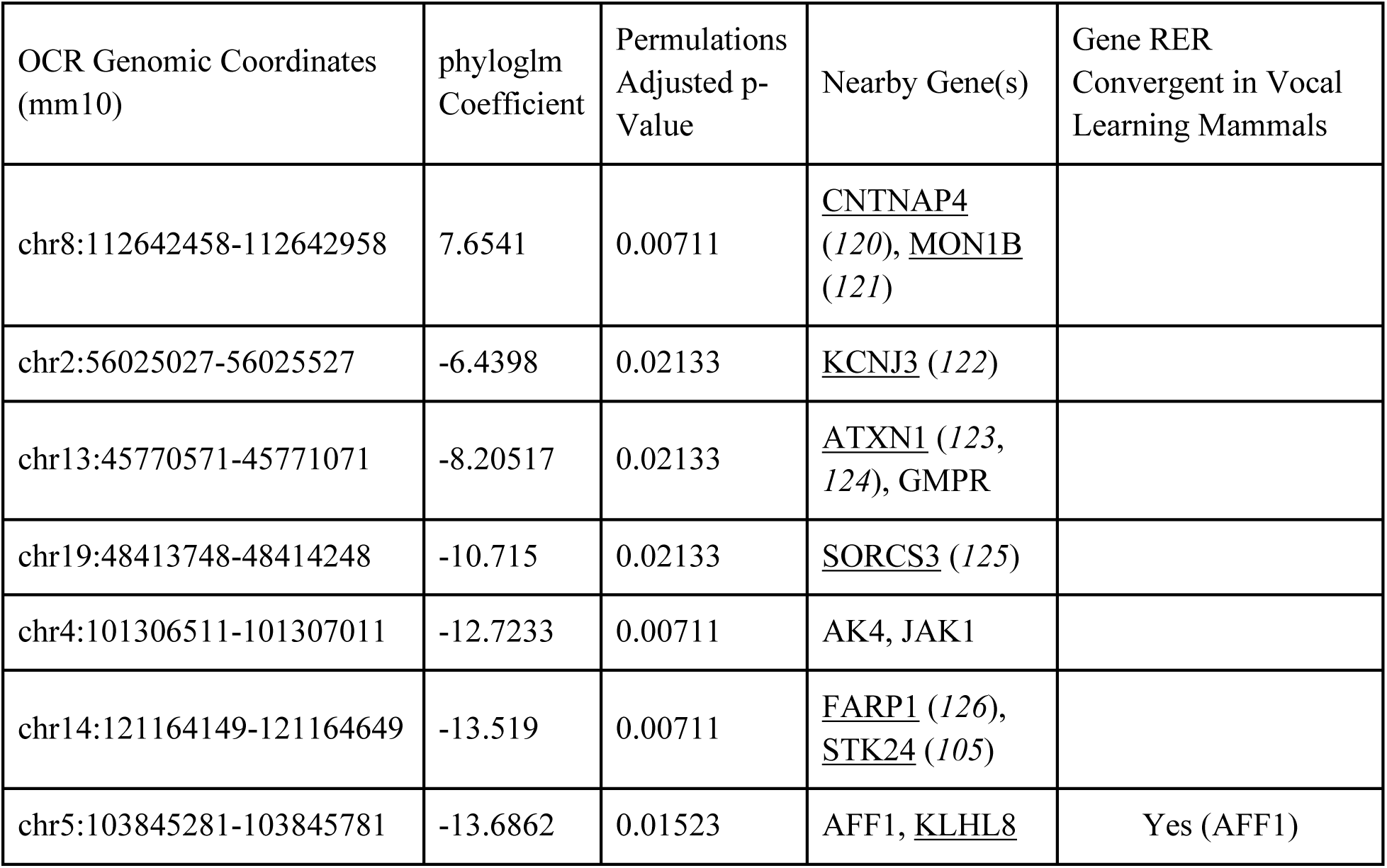
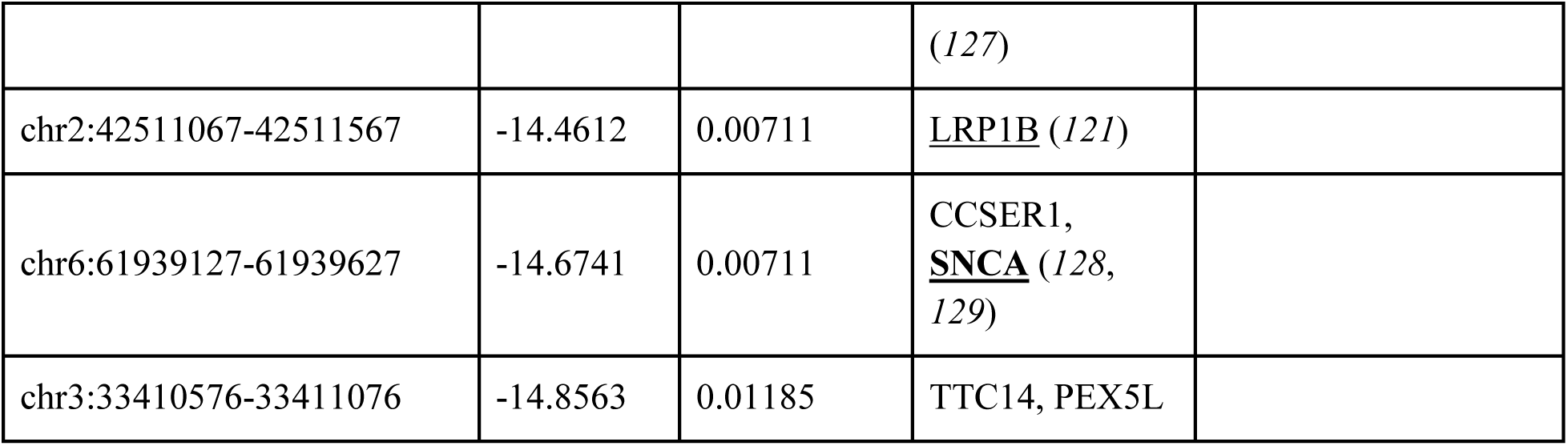
M1 OCRs whose predicted open chromatin state in boreoeutherian mammals is significantly associated with vocal learning. Coordinates are provided for the mouse (mm10) OCR orthologs; otherwise this table is in the same format as Table S1.

## Supplemental Data (separate files)

### Supplemental Data (separate files) Data_S1_VL_RERconverge_gene_sets.xlsx

This file contains a table of results from the RERconverge analysis to identify genes under convergent evolutionary acceleration in vocal learning boreoeutherian mammals, described in the Materials and Methods section “Evaluating the relationship between relative evolutionary rate of protein-coding genes and vocal learning.” For each tab, the first 5 columns provide the raw output of the RERconverge analysis, as described in (*16*). In the first tab (Data_S1A_VL_all_consensus), these values correspond to the analysis considering all 4 boreueutherian vocal learning clades (bats, humans, pinnipeds, and cetaceans). This tab also contains a set of tabs 6 – 9 which indicate with a “1” those genes which were significantly accelerated in each of the subtree analyses, where one of the four vocal learning clades was specifically dropped out from the analysis. The proceeding tabs (Data S1B – S1E) present the output of those analyses. For convenience, the first tab additionally provides columns indicating which of the dropout-robust genes (rows 2 - 202) were also genes associated with TACIT vocal learning-associated OCRs (from Tables S1 and S2) or OCRs with differential ATAC-seq between ofM1 and wM1 (from Data S3).

### Data_S2_VL_RERconverge_gene_functional_enrichments.xlsx

This file contains a table of the results from gene functional enrichment analysis performed using g:Profiler on either the full set of genes significantly convergent in relative evolutionary rate in vocal learning mammals (tab DataS2A_gProfiler_full) or the subset of these genes that were robust to individual vocal learning clade dropout subtree analysis (DataS2B_gProfiler_dropoutrobust), prepared as described in the Materials and Methods section “Gene functional enrichment analyses.” The highlighted rows indicate processes associated with gene regulation.

### Data_S3_ofM1_wM1_differential_OCRs.xlsx

The file contains a table of 348 open chromatin regions (OCRs) discovered from ATAC-seq analyses with differential activity between the bat orofacial and wing subregions of primary motor cortex (ofM1 and wM1, respectively). The first four columns provide locations in the *Rousettus aegyptiacus* genome assembly (mRouAeg1.p), with the fourth column presenting the relative position of the ATAC peak summit in the OCR. Columns 5-9 provide output of DEseq2 analysis (*93*). Proximal genes were identified from the mRouAeg1.p assembly gene annotations (*89*). The full list of TACIT vocal learning associated peaks (intersections from which are provided in column 10) can be found in Tables S1 and S2. The full list of genes under RER convergence in vocal learning mammals (intersections from which are provided in column 11) can be found in Data S1.

### Data_S4_OCR_gene_functional_enrichments.xlsx

This file contains the results of the gene functional enrichment analyses performed using GREAT on the OCRs differential between ofM1 and wM1 (provided in Data S3) mapped to the human genome assembly (hg38), prepared as described in the Materials and Methods section “Gene functional enrichment analyses.” The file contains two tabs, one for the enriched terms from GO Biological Processes (Data_S4A_GOBiologicalProcess) and the other for those from the set of GO Molecular Functions (Data_S4B_GOMolecularFunction).

### Data_S5_OCR_TF_motif_enrichments.xlsx

The file contains a table of transcription factor (TF) binding motif enrichments in the set of peaks differentially active between ofM1 and wM1, the loci of which are provided in Data S3. The description of the MEME analysis that produced these results is provided in the Materials and Methods section “Transcription factor binding motif enrichment.” The results are divided between enrichments in peaks differentially active in ofM1 relative to wM1 (tab Data_S5A_ofM1UP_wM1DOWN) and peaks differentially active in wM1 relative to ofM1 (tab Data_S5B_ofM1DOWNwM1UP).

### Data_S6_VL_species_annotations.xlsx

Table of vocal learning annotations across mammals considered in this study, where “1” indicates vocal learner, “0” indicates vocal non-learner, and “NA” indicates species excluded from the analysis. Description of the preparation of the table, including exclusion criteria, provided in the Materials and Methods section “Coding the vocal learning trait.”

## Supplemental Figures

**Supplementary figure 1.**
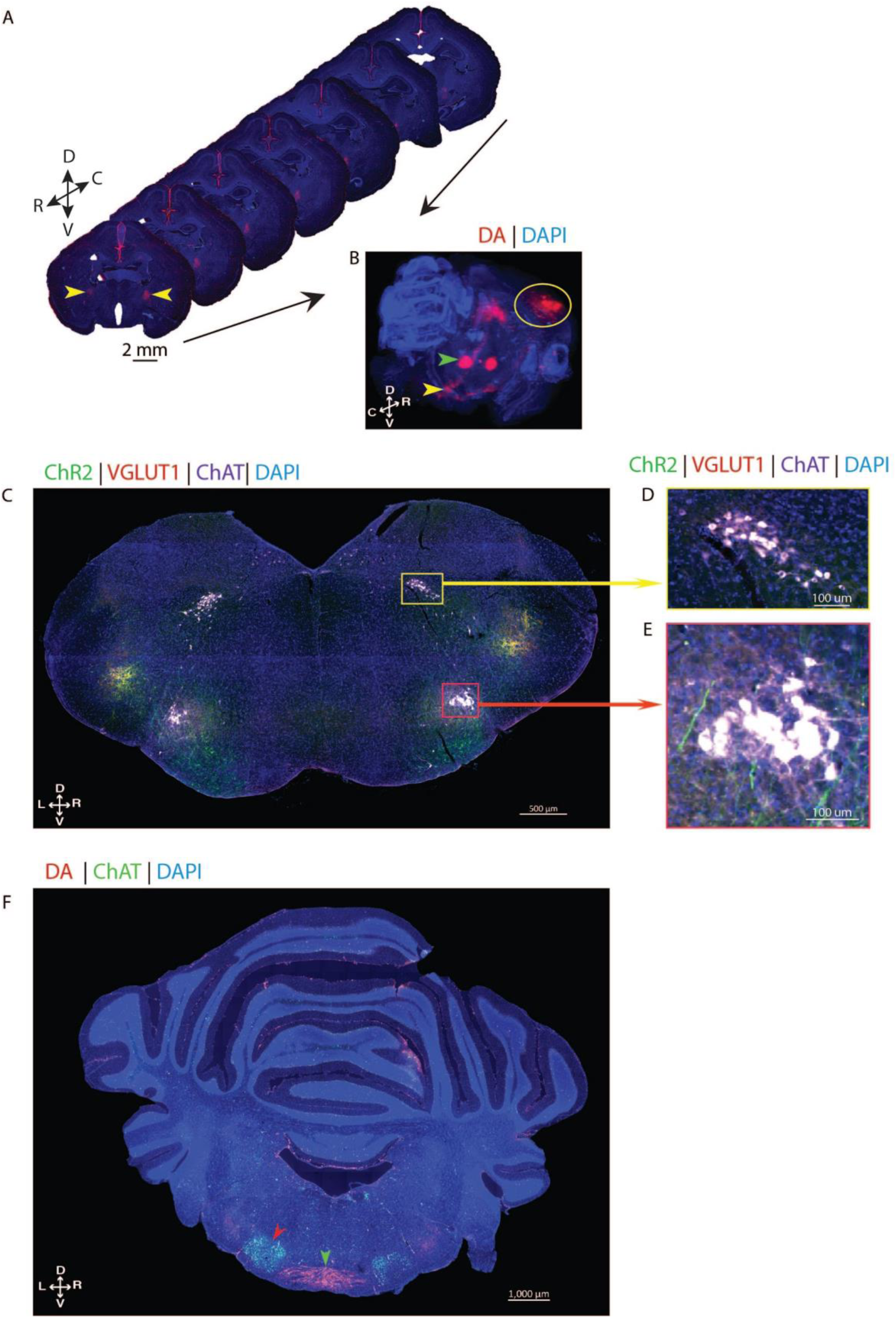
Reconstruction of anterograde and retrograde tracing from orofacial motor cortex (ofM1). **(A)** Example images from one bat injected bilaterally with dextran amine tracer in ofM1 showing seven sequential coronal planes separated by 240 **μ**m, aligned by hand and stacked into a 3D volume shown in (B). The anterograde propagation of the tracer (*red*) is visible in the pyramidal tract (*yellow arrows*) against the background DAPI stain (*blue*). **(B)** 3D side view of the stacked images showing the bilateral anterograde propagation of the tracer (*red*) projecting from ofM1 (*yellow circle*) down the pyramidal tract (*yellow arrow*) to the brainstem. Retrograde propagation of the tracer shows cell bodies in the thalamus (*green arrow*) that send afferents into ofM1. See Supplementary Video 1 for a 3D rotational view of the image stack. **(C-E)** Anterograde tracers are not found in any other brainstem motor nuclei. **(C)** Coronal slice in brainstem following bilateral injections in ofM1 with rAAV5/CamkII-hChR2(H134R)-EYFP (ChR2) labeling fibers in *green*. Synaptic boutons were histologically labeled with VGLUT1 in *red*. Bilateral retrograde injections of CTB (*white*) were delivered within the same bat into the cricothyroid muscles to label laryngeal motoneurons in NA, and in nearby neck and tongue muscles, which enabled the labeling of cells in the hypoglossal nucleus. **(D)** Magnification of hypoglossal nucleus with motoneurons controlling the neck and tongue muscles (*white*). **(E)** Magnification of NA with laryngeal motoneurons (*white*). Note the absence of cortical axons labeled with the anterograde tracer within the hypoglossal nucleus or any other motor nuclei within the medulla except for NA. **(F)** Example coronal section from Egyptian fruit bat medulla showing the decussation of the pyramidal tract (*green arrowhead*) labeled by anterograde injections of fluorescent dextran amine into ofM1. Note the decussation occurs at the level of the facial nucleus (*red arrowhead*), rostral to the location of nucleus ambiguus. Motoneurons are labeled with choline acetyltransferase (ChAT) in *green*.

**Supplementary figure 2.**
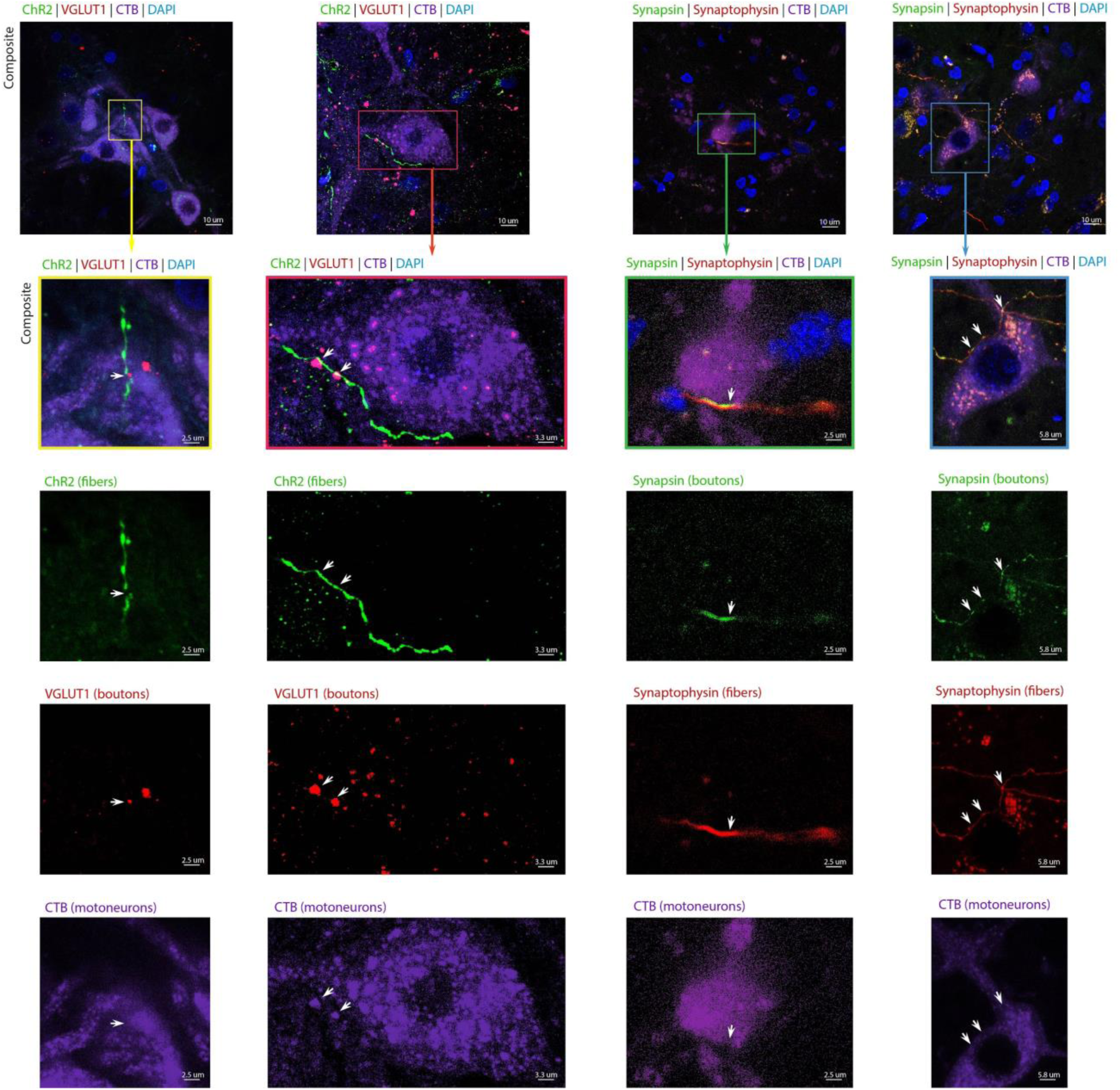
Examples of triple colocalization between corticobulbar fibers, synaptic boutons, and laryngeal motoneurons. (First 2 columns) Example confocal images of nucleus ambiguus (NA) showing triple colocalization (*white arrows*) between corticobulbar axons from ofM1 labeled with rAAV5/CamkII-hChR2(H134R)-EYFP (ChR2, *green*), CTB-labeled cricothyroid motoneurons (*purple*), and VGLUT1-labeled presynaptic boutons (*red*) with a background DAPI stain (*blue*). **(Last 2 columns)** Example confocal image of NA showing triple colocalization (*white arrows*) between corticobulbar axons from ofM1 labeled with AAVDJ-hsyn-mRuby2-T2S-Synap-eGFP (*fibers in red, boutons in green*), and CTB-labeled cricothyroid motoneurons (*purple*), with a background DAPI stain (*blue*). Across all columns, the first two rows are composite images, with the second row being insets of the first row. The three last rows depict three of the four individual channels merged to obtain the insets of the second row.

**Supplementary figure 3.**
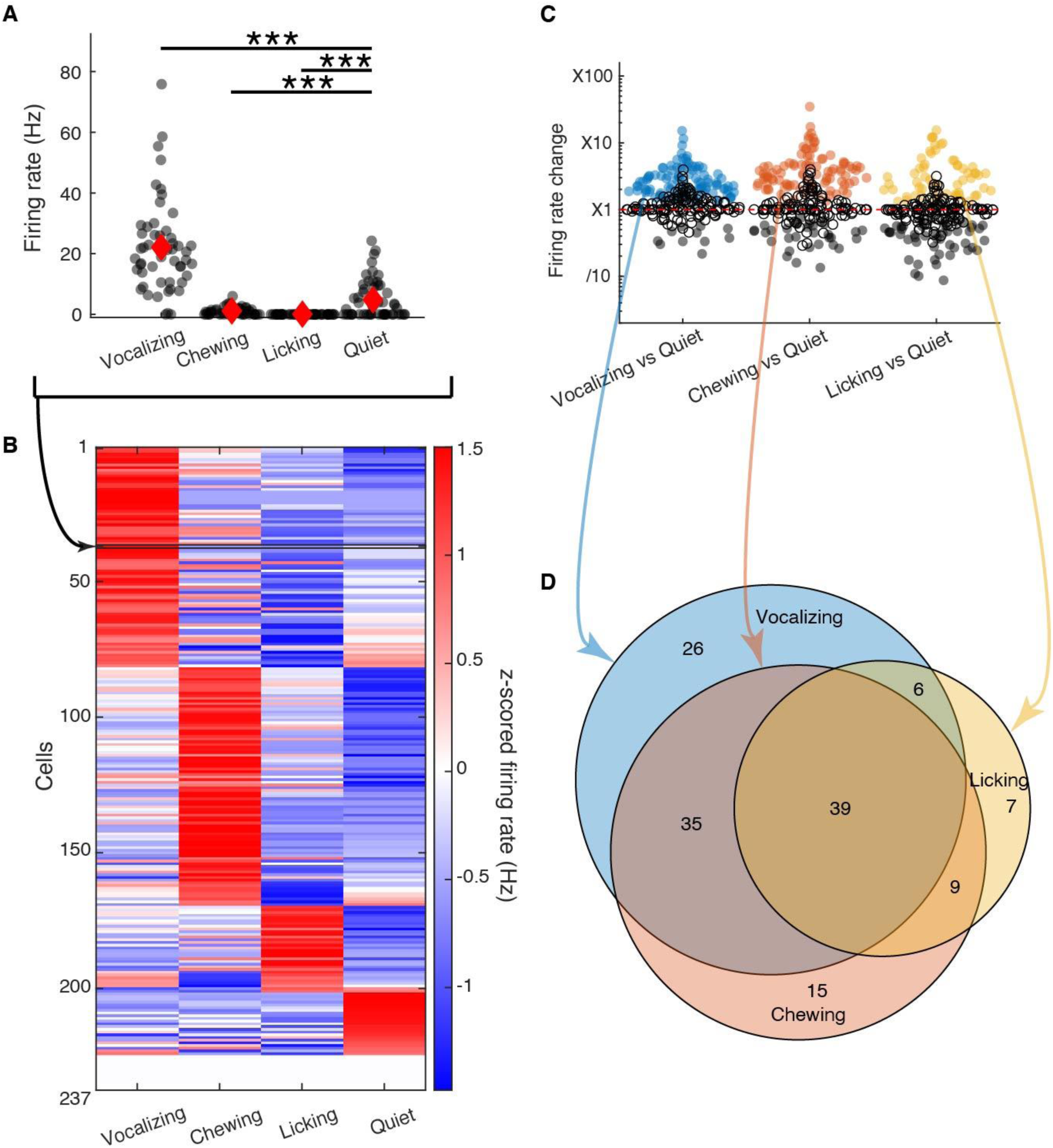
ofM1 neurons activity is modulated by vocalizing, chewing and licking. **(A)** Time average firing rate for an example ofM1 neuron during vocalization production (n=53 vocalizations) and duration matched period of time when the bat was chewing, licking, or quiet and immobile. Black dots are individual events, red diamonds are mean +/- SEM. The neuron was significantly excited during vocal production as compared to quiet and significantly inhibited during chewing and licking as compared to quiet (Anova on Poisson GLM, all p<0.001). **(B)** Z-scored firing rate of ofM1 neurons, the activity of which could be evaluated during vocalizing, chewing, licking, and quiet (*n*=237). **(C)** Change in firing rate during orofacial motor actions as compared to quiet for ofM1 neurons. Each circle represents a neuron. The significance of the Anova on Poisson GLM is signified by filled circles (p<0.001 after FDR correction; *n*=237). As expected, neurons in the orofacial motor cortex are modulated during orofacial motor actions as compared to quiet periods **(D)** Venn Diagram displaying the number of neurons that were excited (significant increase in firing rate as compared to quiet; Anova on Poisson GLM and FDR corrected p-value < 0.001) by vocalizing, chewing, or licking (*n*=137). 26 neurons were excited by vocal production only and not by chewing or licking.

**Movie 1. 3D rotational view of fluorescent dextran amine bilaterally injected in ofM1.** See legend of Supplementary Figure 1A-B for details. The anterograde propagation of the tracer dextran amin (*red*) is visible from the injection point to the pyramidal tract and down to the medulla. A background DAPI stain (blue) is used to better visualize the brain. Note that the decussation of the pyramidal tract, as further depicted in Supplementary Figure 1F, is rostral to the location of NA.

## Notes

### Competing Interest Statement

The authors have declared no competing interest.

